# Pathogen to commensal: longitudinal within-host population dynamics, evolution, and adaptation during a chronic >16-year *Burkholderia pseudomallei* infection

**DOI:** 10.1101/552109

**Authors:** Talima Pearson, Jason W. Sahl, Crystal M. Hepp, Karthik Handady, Heidie Hornstra, Adam J. Vazquez, Erik Settles, Mark Mayo, Mirjam Kaestli, Charles H. D. Williamson, Erin P. Price, Derek S. Sarovich, James M. Cook, Spenser R. Wolken, Richard A. Bowen, Apichai Tuanyok, Jeffrey T. Foster, Kevin P. Drees, Timothy J. Kidd, Scott C. Bell, Bart J. Currie, Paul Keim

**Author notes:** Current Address: GeneCology Research Centre and Sunshine Coast Health Institute, University of the Sunshine Coast, Queensland, Australia (Erin P. Price, Derek S. Sarovich). Current Address: Department of Infectious Diseases & Immunology, Emerging Pathogens Institute, University of Florida, Florida, USA. Current Address: Department of Molecular, Cellular and Biomedical Sciences, University of New Hampshire, Durham, NH 03824.

## Abstract

Although acute melioidosis is the most common outcome *of Burkholderia pseudomallei* infection, we have documented a case, P314, where the disease severity lessened with time, and the pathogen evolved towards a commensal relationship with the host. In the current study, we used whole-genome sequencing to monitor this chronic infection to better understand *B. pseudomallei* persistence in P314’s sputum despite intense and repeated therapeutic regimens. We collected and sequenced 118 *B. pseudomallei* isolates from P314’s airways over a >16-year period, and also sampled the patient’s home environment, recovering six closely related *B. pseudomallei* isolates from water. Using comparative genomics, we identified 126 SNPs in the core genome of the 124 isolates or 162 SNPs/indels when the accessory genome was included. The core SNPs were used to construct a phylogenetic tree, which demonstrated a close relationship between environmental and clinical isolates and detailed within-host evolutionary patterns. The phylogeny had little homoplasy, consistent with a single clonal population. Repeated sampling revealed evidence of genetic diversification, but frequent extinctions left only one successful lineage through the first four years and two lineages after that, resulting in a highly linear topology. Although these extinctions and persistence could be explained by genetic drift, we observe phenotypic changes consistent with *in situ* adaptation. Using a mouse model, P314 isolates caused greatly reduced morbidity and mortality compared to the environmental isolates. Additionally, potentially adaptive phenotypes changed with time and included differences in the O-antigen, capsular polysaccharide, motility, and colony morphology. The >13-year co-existence of two long-lived lineages presents interesting hypotheses that can be tested in future studies to provide additional insights into selective pressures, niche differentiation, and microbial adaptation. This unusual melioidosis case presents a rare example of the evolutionary progression to commensalism by a highly virulent pathogen within a single human host.

**Author summary:** Pathogens frequently jump between different hosts and associated adaptation may lead to the emergence of new infectious agents. Such host-jumping evolution is witnessed through endpoint analyses but these cannot capture genetic changes in lineages that have gone extinct. In this study, we have identified and monitored an example of the adaptive evolution of a bacterium often deadly to its mammalian host, in an unprecedented case whereby disease lessened through time and the pathogen adapted to become a part of the commensal human flora. We used genomic analyses to characterize more than 16 years of this evolutionary process and the stepwise mutations that control dictate pathogen interactions with the patient. Soon after infection, mutational changes occurred that allowed the bacterium to remain in the airways without causing disease. This shift towards avirulence was determined based on clinical data and virulence testing in an animal model. In addition, mutations occurred that contributed to the persistence of the bacteria in the patient’s lungs. Finally, we found evidence for the evolutionary emergence and persistence of two distinct lineages of the bacterium over the last 13 years, presenting interesting questions about niche utilization and separation. Bacteria are ubiquitous in the human body and almost all are beneficial or benign. In this study, we document the evolutionary conversion of a normally deadly bacterium into a commensal.

## Introduction

Within-host evolution of bacterial populations over the course of chronic infections allows the rare opportunity to witness the emergence of bacterial fitness strategies in a new environment. These tactics to evade and overcome host immune responses in order to persist in the host environment may include increased replication efficiency, slower growth, attenuation of virulence, increased transmissibility, and the development of antimicrobial resistance [1]. Genetic diversity within the pathogen population is critical for adaptive evolution as it provides the genetic foundation upon which selection can act. Diversity in an infection may be due to heterogeneity in the inoculating population, lateral gene transfer, or may develop through intrinsic mutations over the course of the infection. The short-term fate of mutations depends on a range of factors such as the population size, environmental stability [2], and the fitness effects of these and other linked mutations. The long-term evolutionary persistence of mutations that emerge within a host depends both on host longevity and pathogen transmission to additional hosts. In most cases, little is known about the emergence of non-fixed mutations that occur during the process of host adaptation. Occasionally, this evolutionary process leads to the emergence of novel species, but only successful lineages are studied and processes are retrospectively inferred. The low cost and high throughput of current sequencing technologies makes characterizing many isolates from single infections possible, enabling a more representative characterization of the population. Phylogenetic analyses of an infecting population and comparison to a close outgroup is necessary to determine the directionality of evolution and to provide a detailed understanding of the population dynamics, evolution, and adaptation.

Melioidosis is an infectious disease that primarily occurs in the tropics, particularly Southeast Asia and northern Australia, where it is hyperendemic [3–5]. Human infections with the melioidosis bacterium, *Burkholderia pseudomallei*, are not usually communicable, with most cases acquired via percutaneous inoculation, inhalation, or ingestion of contaminated soil and water [6]. The incubation period for acute disease following an infecting event is ~9 days, but can be as short as 24 hours with high-level exposure, such as aspiration following near-drowning [7]. Approximately 9% of cases present as a chronic infection, defined as symptoms being present for >2 months [8]. Melioidosis presentations are extremely diverse, ranging from self-resolving skin ulcers without evidence of sepsis to rapidly progressive septic shock with multiple internal organ abscesses [8]. Pneumonia is present in half of cases, overall bacteremia rates are 60%, and septic shock occurs in 21% of all cases [8]. Treatment involves intensive intravenous antibiotic therapy with ceftazidime, meropenem, or imipenem for at least 10 days, followed by eradication therapy with oral trimethoprim-sulfamethoxazole for 3–6 months [7]. Despite this regimen, recurrence occurs in up to 6% of cases through either relapse or reinfection.

Bacterial airway infections are serious illnesses that can progress to chronic infection if not eradicated in a timely manner, leading to morbidity and even mortality. A notable example is the cystic fibrosis (CF) airways, which contain bacterial populations that frequently adapt to evade host and treatment onslaught, enabling pathogens to undergo longitudinal genetic changes *in vivo* that facilitate persistence and drive respiratory decline [9, 10]. Although melioidosis remains rare in CF cases, ~76% of the globally documented cases have led to chronic disease [11] compared with just 10% in non-CF cases. *B. pseudomallei* can undergo dramatic genomic and transcriptomic changes that may enable persistence in the airways of CF patients despite intensive antibiotic treatment regimens [12, 13], demonstrating that this bacterium is adept at long-term airway persistence under certain conditions. In contrast, acute pneumonia can be fulminant with subsequent sepsis. Melioidosis is notable for having both scenarios in the clinical spectrum [14]. The progression of inhalation/pulmonary melioidosis can be very rapid and severe. In animal models, acute melioidosis occurs within 1–2 days of aerosol exposure to *B. pseudomallei* [15].

Similar to many other saprophytic soil dwelling bacteria, *B. pseudomallei* hosts a large and highly plastic genome [16] that facilitates rapid adaptation to dynamic and diverse environmental challenges. The genome of wild-type *B. pseudomallei* isolates is large (~7 Mbp) and is carried on two chromosomes. However, the size of the pangenome carried by multiple conspecific strains is truly impressive. The accumulation of coding region sequences in the pan-genome [17] is largely associated with the horizontal acquisition of sequences, including genomic islands from genetic near neighbors [18]. The virulence of *B. pseudomallei* has been attributed to multiple genomic loci, including type III and type VI secretion systems, lipopolysaccharides, actin-based motility factors, toxins, adhesins, and capsular polysaccharide clusters [16, 19]. The large accessory genome likely contributes to the observed clinical diversity [20] and results in the ability to rapidly adapt to dynamic and hostile environments.

In the Darwin Prospective Melioidosis Study (DPMS), 9% of the 1,105 culture-confirmed cases over the last 29 years presented as a chronic infection [8]. The majority of patients in these cases have survived, and all but one eventually cleared their infection. This individual, Patient 314 (P314), remains sputum culture-positive for *B. pseudomallei* since being diagnosed with melioidosis in 2000 as an in-patient following a bronchiectasis exacerbation event [21]. We have collected and cultured *B. pseudomallei* isolates from P314’s airways since this initial diagnosis to better understand the population dynamics in this unprecedented case. We have previously shown genetic diversity in variable-number tandem repeat loci among these isolates [22] and subsequently compared three P314-derived genomes to document some of the evolutionary changes over an 11.5-year period [21]. Here, we analyze and compare the genomes and phenotypic characteristics of 118 *B. pseudomallei* isolates sampled over a >16-year chronic carriage period from P314’s airways, along with six closely-related isolates sampled from P314’s home between two and four years after diagnosis. These comparisons allow assessment of the evolutionary and temporal dynamics of this extraordinary case rather than strictly comparing evolutionary end-points, providing broader inferences on the evolution of host specificity.

## Materials and methods

### Clinical sample collection, culturing, and isolation

We collected and analyzed a >16-year longitudinal series of *B. pseudomallei* isolates for this study. The Microbiology Departments of the Royal Darwin Hospital, Darwin and The Prince Charles Hospital, Brisbane, and the Melioidosis laboratory at the Menzies School of Health Research, Darwin, performed bacterial isolations. *B. pseudomallei* was cultured from sputum, nasal and throat swabs, nasal mucus, or extracted lung tissue following a lobectomy in 2003 by using Ashdown’s selective broth and Ashdown’s selective medium [23], except for the initial clinical isolate MSHR1043, which was isolated using chocolate agar as it is sensitive to gentamicin [21]. *B. pseudomallei* was confirmed biochemically using the API 20E system (API System SA, Lyon, France) or, more recently, MicroScan WalkAway (Dade Diagnostics, West Sacramento, Calif.) and the TTS1 *B. pseudomallei* PCR assay [24].

### Genome sequencing, assembly, and data deposition

Genomes were subjected to whole-genome sequencing (WGS) on the Illumina platform, followed by assembly with SPAdes v3.10.0 [25]; contigs that aligned to non-target species (e.g. PhiX) or that contained anomalously low coverage compared to the average genome coverage were manually removed. The environmental isolate MSHR1435 was sequenced on both the Illumina and PacBio platforms and assembled as described elsewhere [26]. This closed genome was previously deposited into GenBank under accessions CP025264.1 and CP025265.1. The raw read data for each strain were submitted to the Sequence Read Archive database (Table S1) under BioProject accession PRJNA321854.

### SNP calling

Short reads from all genomes were mapped across the complete genome of MSHR1435 using BWA-MEM v0.7.7 [27], and SNPs were called with the UnifiedGenotyper method in GATK v3.4.6 GATK [28, 29]. SNPs were filtered from downstream analyses if they fell within duplicated regions in the reference genome, as determined by a NUCmer self-alignment [30], if they fell within short tandem repeats, as determined by Tandem Repeats Finder v4.0.9 [31], or if they had a coverage depth of <3 or an allele proportion <90%. All of these functions were wrapped within the NASP pipeline [32]. Corresponding SNPs between different reference genomes (MSHR1435, MSHR1043, K96243) were identified with the ISG pipeline [33]. Functional information was applied to each SNP with SnpEff v4.3r [34] using the PGAP annotation.

### Phylogenetic reconstruction

The phylogenetic tree was constructed in two steps. First, 126 orthologous, core-genome SNPs (S2 Table) present in each isolate (except 2 missing SNP loci in the outgroup K96243, which did not affect the topology) were used to construct a maximum parsimony tree using PAUP 4.0b10 [35] to give an accurate estimation of phylogenetic grouping, albeit with limited resolution. This tree was one of four equally parsimonious trees. Two homoplastic SNPs among clinical strains caused a consistency index (CI) (i.e., excluding uninformative characters) of 0.9737 and a tree length of 128. The tree shown in Fig 4 is the single tree whose topology agrees with a tree inferred without the homoplastic SNPs. Secondly, other characters (indels) were added to the SNP matrix for added phylogenetic resolution to better capture all the character changes that occurred along all branches of the tree. These additional characters also included deleted regions with and without SNPs. To ensure independence of evolutionary events, each inferred deletion event was manually added as a single binary character to the data matrix. Similarly, for SNPs present in regions deleted in other isolates, the inferred immediate ancestral state for each indel was determined (from the phylogenetic placement of the sample in the first tree) and added to the matrix. For SNP calls that were missing in some genomes, we manually determined if the missing call was due to a deletion or poor sequencing coverage. For loci in low coverage areas, we assumed the immediate ancestral allele as a conservative estimation of the evolutionary state. The unambiguous optimization of these character states eliminates potential phylogenetic ambiguity due to missing data in tree reconstruction [36] and treats each deletion event as a single event rather than a separate event associated for every SNP within that region. This method is similar to uninode coding [37]. Sequences for samples MSHR1360 and MSHR1459 are from a population of cells with two discrete genotypes, which were not parsed during the subculturing step. For some SNPs in these sequences, both alleles are detectable and the deleted regions are present in a minority of reads, although each allele had a minimum coverage of 3x. Based on the presence of alleles, we could determine the phylogenetic position of both components and therefore include them in the final tree as they contribute to our understanding of the population dynamics. Other sequences also show evidence of fine-scale mixed genotypes, however minor alleles occur in very few reads and we could not confidently parse the genotypes.

### Comparative genomics

To identify deletions, raw data were aligned against MSHR1435 with BWA-MEM and the per base depth of coverage (DOC) was identified with SAMtools v0.1.19 [38]. Due to low levels of genome mixtures observed across many of the genomes, a dynamic filtering approach was employed. First, regions with <25% DOC compared with the average DOC for each genome was identified by summing the coverage across each base of the reference genome and then dividing by the total genome length. For each base, if the depth was less than 25% of the average DOC, then the position was were considered absent. Visual confirmation of deletions was performed using Circos [39]. To further verify deletions, each coding region sequence (CDS) in MSHR1435 was screened against all genome assemblies with the large-scale BLAST score ratio (LS-BSR) pipeline [40] using the BLAT aligner [41]. CDSs that fell within deletions identified with the read mapping were then tabulated (S3 Table). LS-BSR was also used to find CDSs that were part of the core or the accessory genome. For one region that was present in environmental samples but had signatures of degradation in clinical samples, as identified with LS-BSR, raw data were re-aligned to MSHR1435 using Bowtie2, and SAMtools was used to identify per base DOC. This approach allowed for the determination of deletion boundaries.

### Molecular Clock Analysis

To reconstruct the evolutionary history of the P314 clinical isolates while simultaneously estimating the rate of intra-patient evolution and time to most recent common ancestor for clinically relevant events, we performed a Bayesian molecular clock analysis in Bayesian Evolutionary Analysis Sampling Trees (BEAST) v1.8.4 [42] using the 126 core-genome, orthologous SNP dataset. Substitution model selection was carried out in MEGA 7.0.9 [43] for the 120 clinical genomes. The corrected Akaike Information Criterion and Bayesian Criterion results indicated that the HKY substitution model [44] best fit this dataset. To determine the best fitting clock and demographic model combinations for these data, we employed the generalized stepping-stone sampling marginal likelihood estimator [45]. Each of the eight model combinations was iterated 100,000,000 times, where Markov chains were sampled every 10,000 generations. We found that the strict clock combined with the Bayesian Skygrid demographic prior outperformed the other seven combinations (Table S4). Using the Strict Skygrid model combination, we ran three additional chains for 100,000,000 generations, sampling every 10,000, and found convergence within and among chains using Tracer v1.6 [46]. Importantly, we included a correction for SNP ascertainment bias by indicating the number of invariant sites (A: 985225, C: 2103022, G: 2112938, T: 982967), allowing for a more accurate estimation of rates and timing. We used LogCombiner [42] to merge the four different chains, discarding the first 10% as burn in (40,000,000 generations), and then resampled every 40,000 generations. The resulting file was input to TreeAnnotator to produce a maximum clade credibility tree, and then visualized using FigTree v1.4.3 [47].

### Phenotypic testing

We previously noted considerable differences in growth rate and colony morphology between P314’s initial isolate and a 139-month isolate [21]. To determine if other isolates over the course of the infection also had observable changes in growth characteristics, 110 P314 isolates were streaked for isolation onto modified Ashdown’s agar [23] minus gentamicin. Plates were incubated at 37°C. At ~48 h and again at 72 h, plates were photographed. Photographs were used to examine and compare growth characteristics and to determine colony morphology among all isolates. Colony morphology was mapped onto the phylogenetic tree.

### Motility testing

Flagellar motility is a virulence factor that can be recognized by Toll- and NOD-like receptors [48]. Motility is easily tested in the laboratory by stab-inoculating a semi-solid media containing a bacterial growth indicator; if bacteria are motile then growth will be observed diffusing from the stab line. After ~48 h incubation at 37°C on Ashdown’s agar (minus gentamicin), single colonies or cells from a visible region of uniform growth were selected and stab-inoculated into motility test medium with tetrazolium growth indicator (Hardy Diagnostics, California). A known motility-positive strain of *B. pseudomallei* unrelated to P314, Bp82, and *Staphylococcus aureus* were used as motility-positive and -negative controls, respectively. Controls were grown on standard Luria Bertani agar for ~48 h before use. Additionally, a sterile inoculating needle was used to stab a single vial of motility medium and used as an uninoculated control. Motility tests were incubated at 37°C; tubes were observed for growth at 48, 72, and 120 h and determined as positive or negative for motility based on comparison to the controls [49]. Results were mapped onto the phylogenetic tree.

### Capsular Polysaccharide (CPS) testing

Previously identified mutations in *wcbR* in the P314 isolates [21] probably led to decreased virulence of these isolates by reducing CPS-I production and expression, an essential virulence factor [50]. To test this hypothesis, we examined the CPS production using the InBiOS Active Melioidosis Detect™ Rapid Test (Seattle, WA), which specifically detects CPS produced by *B. pseudomallei* and *B. mallei* [51]. Tests were performed using the 110 ~48 h-old isolates described above and according to the manufacturer’s instructions. Additionally, CPS and lipopolysaccharide (LPS) production was evaluated in seven isolates by Western blot analysis.

### Antibiotic testing

Certain P314 isolates were screened using E-tests to determine their minimum inhibitory concentrations (MICs) to assorted clinically relevant antibiotics, including gentamicin, ceftazidime, doxycycline, meropenem, amoxicillin-clavulanate, and imipenem. For each antibiotic, test strips were placed on bacterial lawns and plates were incubated and interpreted as per manufacturer’s instructions (bioMérieux, France).

### Animal testing

Mouse challenge experiments were performed under Select Agent and ABSL-3 containment practices. This work was conducted at Colorado State University and was performed in strict accordance with the recommendations in the Guide for the Care and Use of Laboratory Animals of the National Institutes of Health. Every effort was made to minimize animal suffering. The protocol was approved by the Animal Care and Use Committee of Colorado State University (protocol number 17-7497A). BALB/c mice were purchased from Charles River laboratories and housed at Colorado State University. To examine the attenuation of the P314 and P314-associated environmental isolates, isolates were grown at 37°C in BHI medium to an OD_600_ of 1, supplemented with glycerol to 10% v/v, and frozen in multiple aliquots at −80°C. A vial of these stocks was thawed and serial dilutions plated on BHI agar plates to determine titer. Mice were intranasally challenged as previously described [52]. The amount of bacterial culture (CFU/dose) was verified by titration and plating the challenge inoculum on BHI agar plates. Mice were weighed and evaluated for clinical symptoms associated with *B. pseudomallei* infection prior to and after infection. This monitoring was performed daily for the first 7 days, and subsequently every 3–4 days to 21 days. The clinical scores indicated in the study were: 0 = normal, 1 = questionable illness, 2 = mild but definitive illness, 3 = moderate illness, 4 = severe illness, moribund – euthanized, 5 = found dead.

### Human Subjects Ethics Statement

This study involved the informed consent of a single human subject (P314), with ethics approval obtained as previously described [21].

## Results

### A chronically infected patient

Melioidosis occurs in an uneven distribution across northern Australia, with the pathogen found in soil and water where cases occur [53]. Acute melioidosis dominates the clinical presentations and chronic infections that fail to resolve despite appropriate antimicrobial therapy, leading to long-term carriage, are very uncommon. Apart from the case presented here, such cases have only been recognized in a very small number of melioidosis patients who have cystic fibrosis [11, 13]. In July 2000, a 61-year-old female patient (P314) with known bilateral non-cystic fibrosis bronchiectasis and past respiratory infections with *Mycobacterium avium* complex and *Pseudomonas aeruginosa*, presented with worsening cough and infiltrates on her chest X-ray consistent with a diagnosis of acute pneumonia. *B. pseudomallei* was cultured from her sputum as well as nose and throat swabs, confirming her first presentation of melioidosis. She was initially treated with 14 days of intravenous ceftazidime and her symptoms improved, but her sputum continued to culture *B. pseudomallei*. Subsequent multiple courses of antibiotics in various combinations over the next 10 years failed to clear her infection. These included intravenous ceftazidime, intravenous meropenem, nebulized ceftazidime, oral doxycycline and oral amoxicillin/clavulanate. She has a severe allergy to trimethoprim/sulfamethoxazole, which precluded its use. Respiratory symptoms and quantity of productive sputum production have fluctuated over subsequent years, but both improved substantially after she had a left lower lobectomy in August 2003, 37 months after initial diagnosis (Fig 1). Nevertheless, despite being less symptomatic than initially, she still has daily production of sputum and remains *B. pseudomallei* sputum culture-positive to this day, but has not been on antibiotics with activity against *B. pseudomallei* for much of the last five years. We have collected and analyzed a >16-year longitudinal series of *B. pseudomallei* isolates for this study.

**Fig 1.**
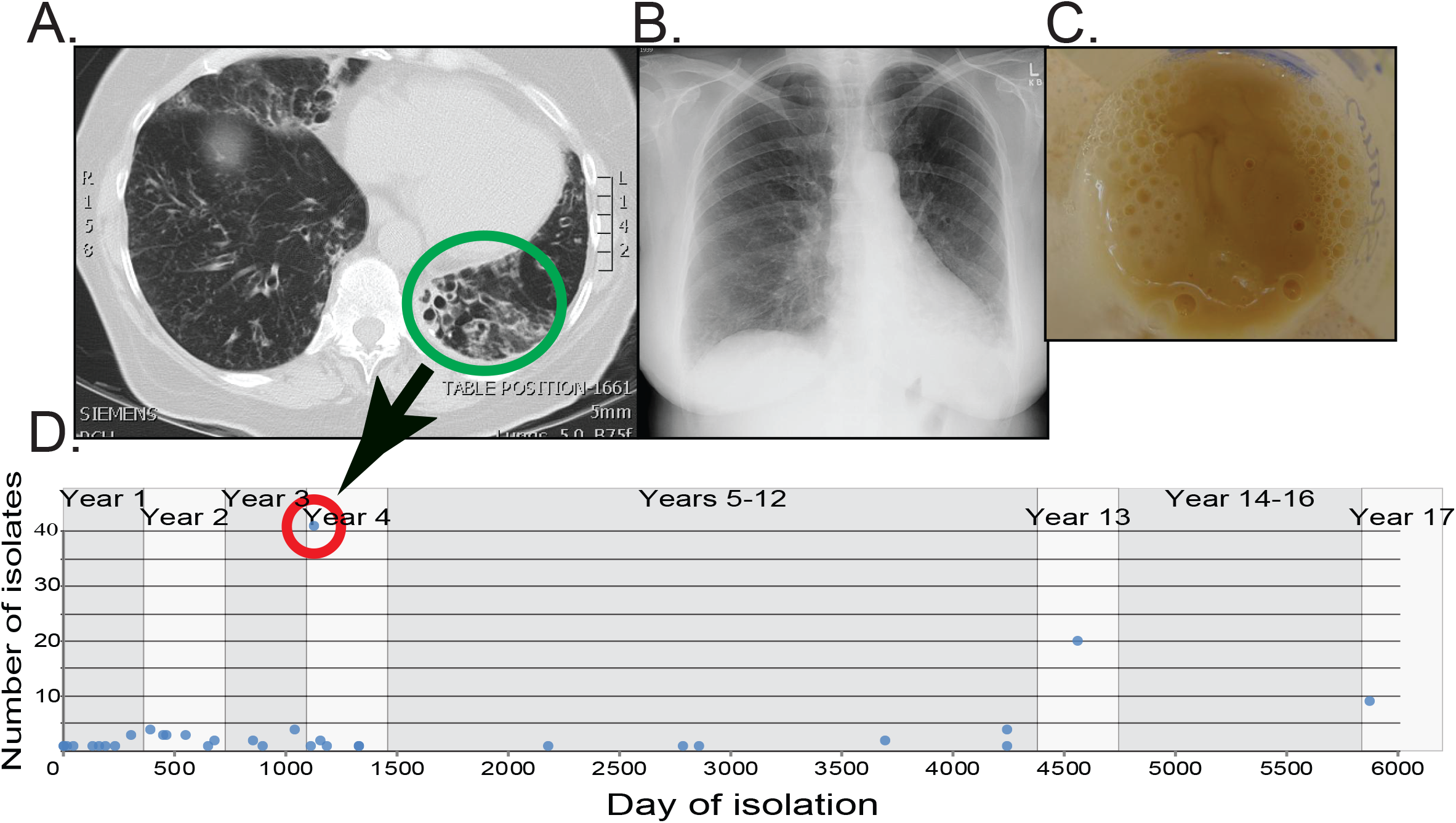
Patient Timeline of Treatment and *B. pseudomallei* Isolate Collection. A) CT scan shows damaged, widened airways (green circle) in the lower left lobe that required surgery on day 1134. B) Chest X-ray after lobectomy. C) Patient’s sputum in cup. D) Timeline of isolate collection. From excised lung tissue, we cultured 41 isolates on day 1134 (red circle).

### Environmental Isolates

To identify the putative origin of her infection, P314’s home environment was repeatedly sampled over an approximate two-year period (Aug 2002, Oct 2002, Jan 2003, May 2003 and Apr 2004). At the time, P314 lived in a semi-rural town in the tropical north region of the Northern Territory, Australia, and her home used unchlorinated groundwater from a private well for consumption and bathing. *B. pseudomallei* was cultured from the bore head (water prior to the tank), holding tank, pool, and household delivery pipes. In all, 21 environmental colonies were isolated and genotyped from 11 samples using multilocus sequencing typing and WGS. Six of the environmental isolates were multilocus sequence type (ST) 131, matching the patient isolates. At the WGS level, all six were highly similar to the patient’s clinical isolates. Genome sequences from the clinical and the six ST131 environmental isolates were compared to three other ST131 genomes (from other patients) to understand within-ST131 diversity (S1 Fig). We found that there were a maximum of 16 core genome SNPs among the different environmental isolates and the closest environmental isolate (from pool water) differed from the closest P314 clinical isolates by only 1 core genome SNP. However, no SNP separated all environmental isolates from the clinical isolates, consistent with probable evolution of environmental isolates from the original infecting ancestor in the ~2 years following the original infection event. In contrast, the phylogenetic distance of the P314-associated strains to ST131 isolates from other clinical cases was much greater. There was an average SNP distance of nearly 10,000, and in some cases 40,000 core genome SNPs, between isolates in this more diverse set. Thus, environmental sampling at P314’s home point strongly towards infection occurring from her home environment and provided strains that closely represent the original infecting agent.

### Genomic Deletions

Seven regions that include large deletions were observed among the clinical isolates when compared to the reference genome of the environmental isolate, MSHR1435. Some of these deletions became fixed in the population, others occurred in lineages that went extinct, and others persisted in a subset of the population. Overlapping deletions with different start and end points, suggesting independent events, occurred in two regions. Subsequent decay surrounding previously deleted regions occurred in two additional regions. These deletions are depicted in Fig 2 while the phylogenetic location of each deletion event can be seen in Fig 3 and S3 Table. On Chromosome 1, we saw only two deleted regions. Deletion 1 (~31,000 bp) was observed in a single genome (MSHR1287) while Deletion 2a (1,416 bp) was identified in all, with the exception of four early clinical genomes. Three subsequent adjacent deletions (2b, 2c, 2d) occurred in a single lineage. All other large deletions were found on Chromosome 2. Deletions in regions 3 and 4 occurred relatively early over the evolutionary history of these genomes and while deletions 3 and 4a became fixed in the population, the 4b deletion event occurred independently on a lineage that subsequently went extinct. Regions found elsewhere in the genome show a high level of similarity to a small part of deletions 3 and 4a, giving the appearance in Fig 2 that each of these deletions are actually comprised of two separate deletions. Mixed-culture genomes (see below) also resulted in low levels of coverage in regions that were expected to be deleted. Examples of this can also be found in deletions 3, 4a, and 6a. Deletions in regions 4, 6, and 7 each involve the elimination of multiple overlapping genome segments, representing independent events and parallel evolution. Only one of these 15 deletions was associated with all clinical isolates, suggesting that this initial deletion may be associated with an important early adaptation and the loss of virulence. This deletion is in the non-ribosomal protein synthetase gene cluster. This is a highly modular enzyme and the loss of this intragenic region may not eliminate its expression, but rather change the secondary products it produces [54].

**Fig 2:**
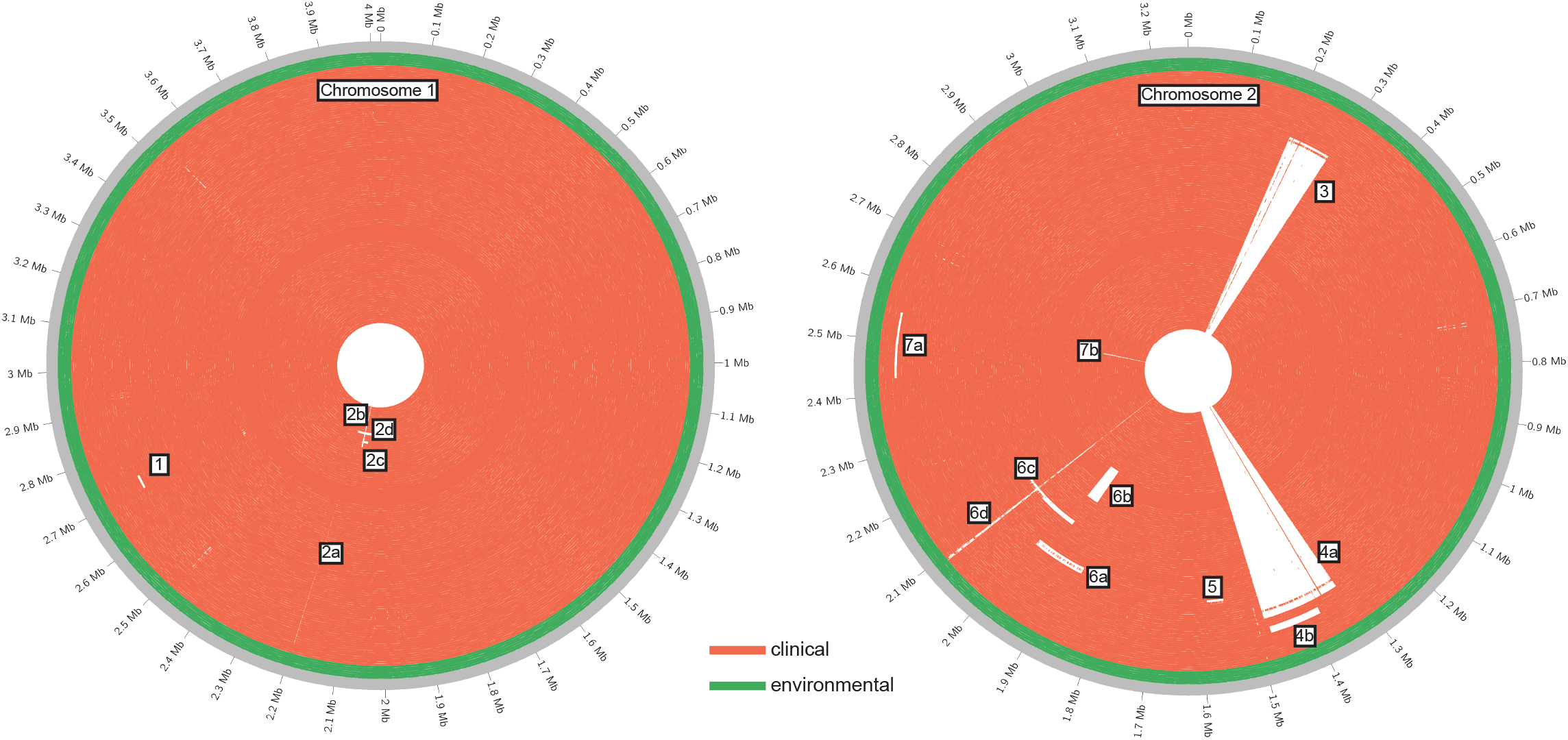
The location of large deletions in P314 genomes relative to MSHR1435. Whole genome sequencing of 120 *B. pseudomallei* collected over >16 years reveals large chromosomal deletions across both chromosomes. The genomes’ order from the outside to inside concentric circles corresponds to the top to bottom order on the phylogenetic tree (Fig 3). This inside-to-out arrangement approximates the evolutionary trajectory of the infecting *B. pseudomallei* population. MSHR1435, which is used as the reference, is outermost and is a closed genome while all others are high quality drafts. All environmental isolate genomes (including MSHR1435) are basal in the phylogeny (Fig 3) and found in the outermost rings (green) of the concentric circles. Raw sequence reads were mapped to the reference chromosomes and any base with a depth of coverage of <25% of the average genome coverage was considered to be missing. The figure was constructed with Circos [39].

**Fig 3.**
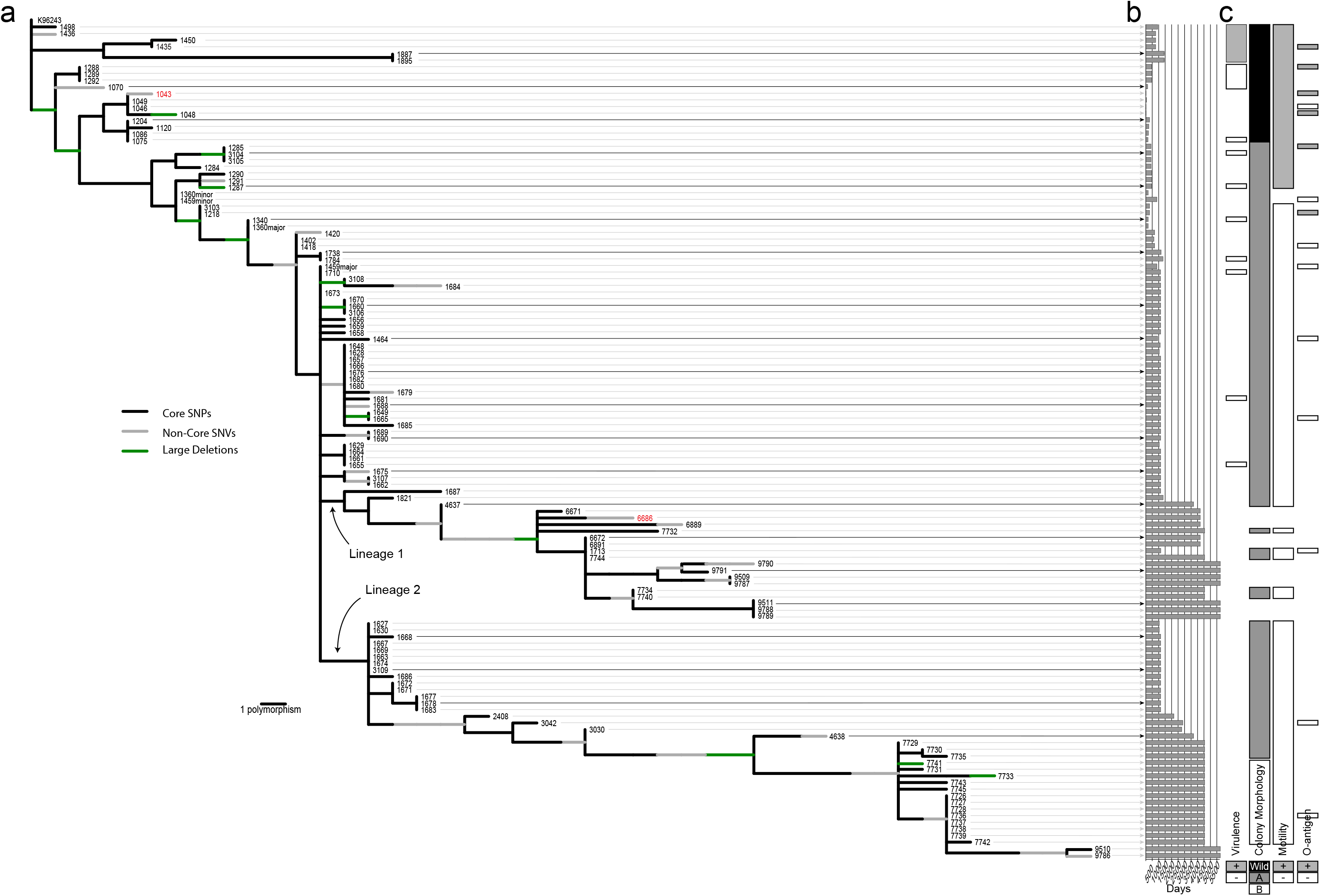
Evolutionary changes. (a) Phylogenetic tree showing evolutionary locations of polymorphism, (b) day of collection for each isolate, and (c) morphological characteristics (virulence, motility, and three colony morphotypes - W= wild type, A = small colony variants, B = fastidious). Core genome SNPs are represented as black branch segments, while non-core genome SNPs are gray. The large deletions shown in Fig 2 are green branch segments. We previously compared the genomes in red [21]. Where mixtures are observed, the major and minor components are labeled.

### Deleted genes

Some genes contained within annotated deletions were associated with known virulence mechanisms, including the type III secretion system (CXQ84_30725-CXQ84-30860) [55]. The single, and earliest, deleted region is found in all clinical isolates, and is thus likely associated with virulence attenuation. This region encompasses an annotated non-ribosomal peptide synthetase (CXQ84_31440), a large (3,290 aa), repetitive gene associated with deletions in regions 6c and 6d (Fig 2). This is shown in higher resolution in Fig 4. AntiSmash [56] results suggest that this region is annotated as a condensation domain-containing protein (SMCOG1127), although its potential product is unknown.

**Fig 4.**
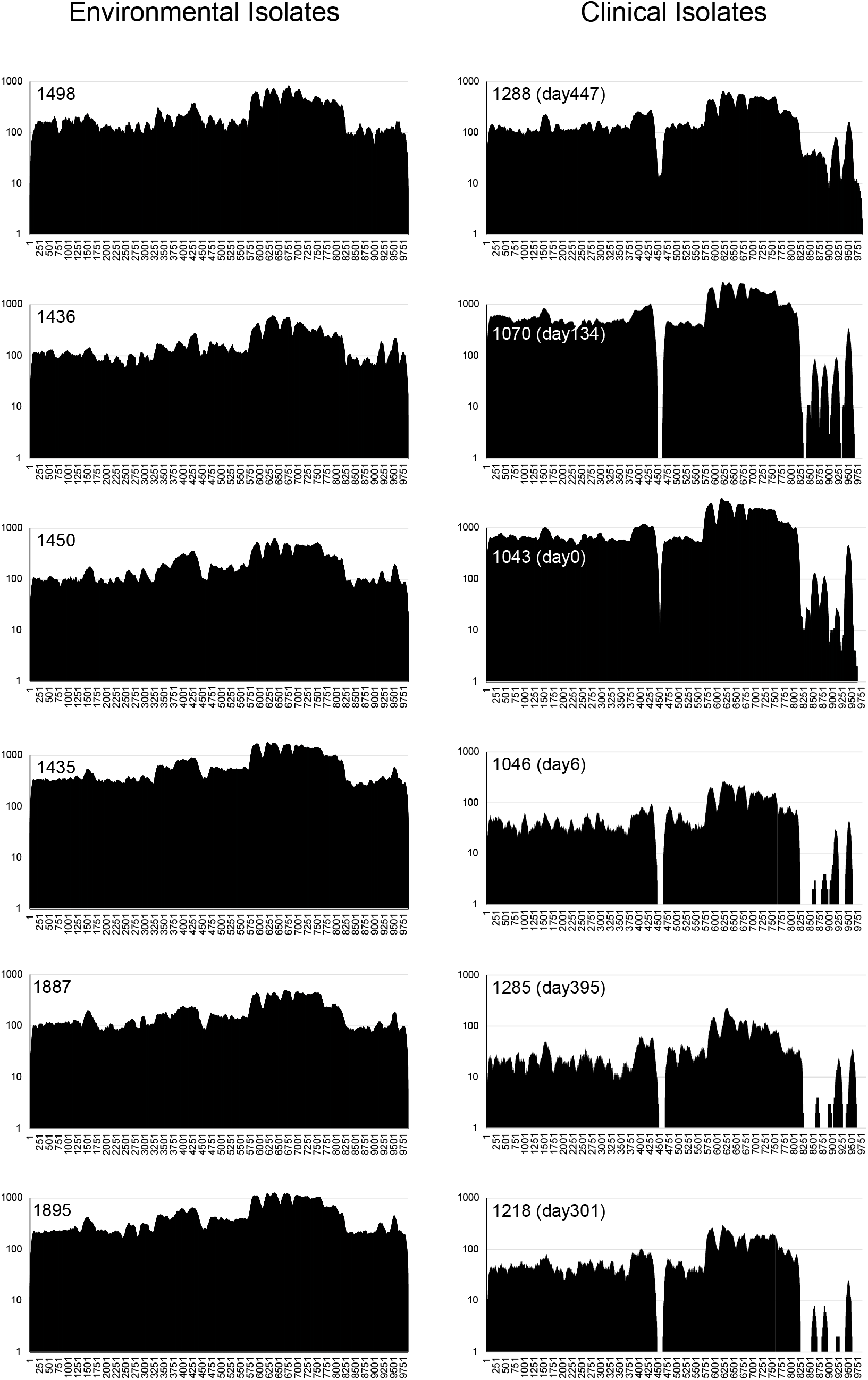
Sequence Read Mapping to a Non-Ribosomal Peptide Synthetase (NRPS) Gene from Different Isolates. The Illumina reads from each of the isolates in this study were aligned to the NRPS gene from the MSHR1435 reference (MSHR1435 locus CXQ84_31440). This figure is a higher resolution representation of the read mapping shown in Fig 2 and covers the 6c and 6d regions shown there. Twelve examples of this analysis are illustrated with six environmental and six clinical isolates, with the complete isolate set present in the supplemental material (S1 Table). The read mapping shows the early loss of the internal gene sequences around position 4500 and later deletions in the 3’ region. Isolate are arranged from top to bottom in the same order as in the phylogenetic tree (Fig 3). These same isolates were all used for animal challenge experiments (Fig 6).

The CDSs from the large deletions shown in Fig 2 were also screened against 51 *B. mallei* genomes (S5 Table) with BLAT in conjunction with LS-BSR. The results (Fig 5, S6 Table) demonstrate that many deletions in P314 have also been observed in *B. mallei*, suggesting parallel evolution, possibility associated with host adaptation in the respiratory tract. Many of these *B. mallei* isolates are still virulent, suggesting that the observed parallel evolution is not necessarily associated with pathogenesis.

**Fig 5.**
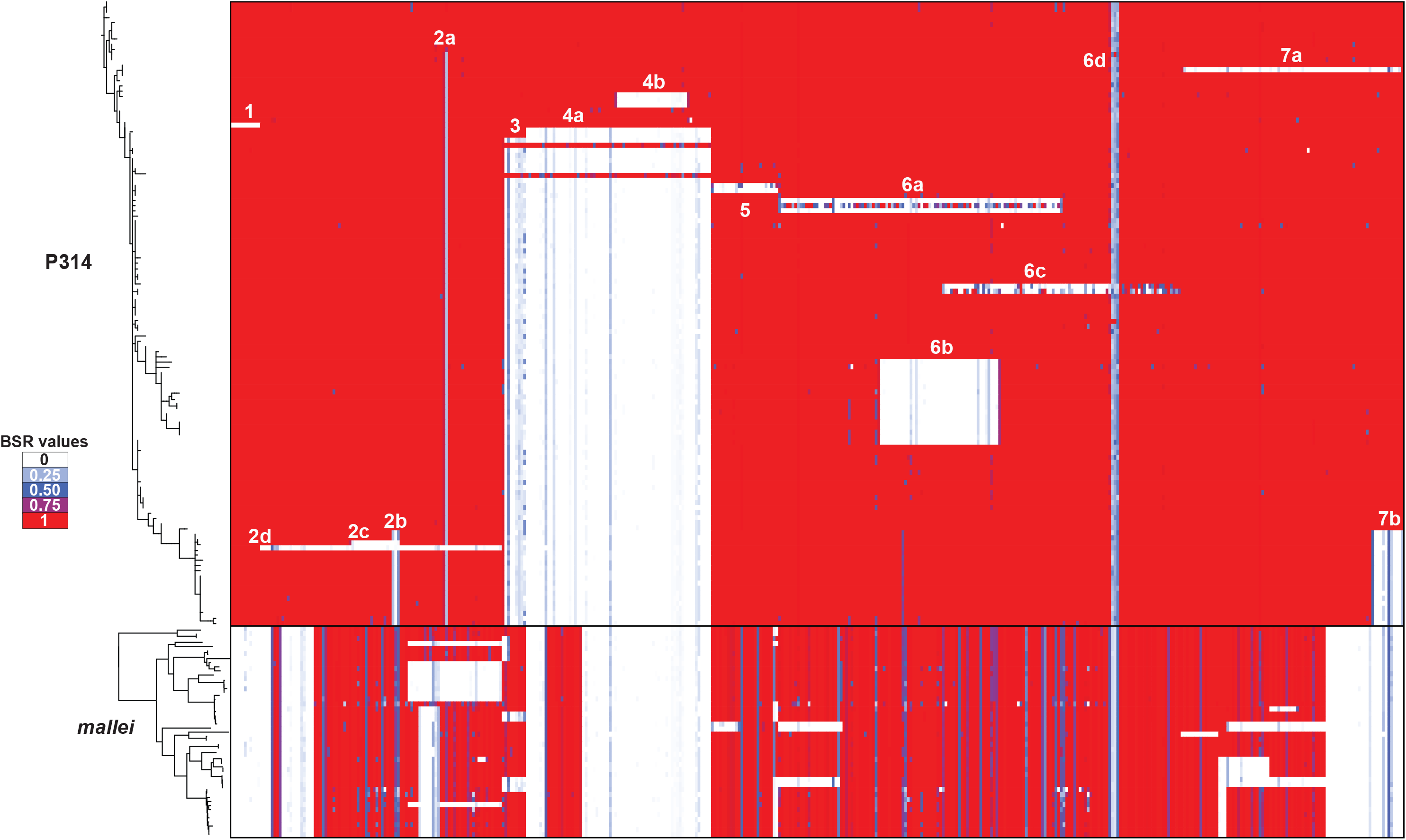
Convergent mutations between *B. mallei* and P314. A phylogeny of P314 and 51 *B. mallei* genomes mapped with the seven genomic deletions (Fig 2) identified among P314 genomes. The heat maps represent only the subset of coding region sequences (CDSs) (n=437 of 815 CDSs) found in the deleted regions. Parts of each of the seven regions deleted among P314 genomes were also deleted in *B. mallei* genomes. Deletions in the 6d region corresponds to the NRPS gene (Fig 4) and are also observed in *B. mallei* (Fig 5).

### Small genomic deletions

We have previously described a small number of indels in P314 genomes [21]. These regions were screened against all P314 genomes and mapped across the phylogeny (S7 Table). The *wcbR* frameshift (CXQ84_04665), which results in an altered capsular polysaccharide synthesis protein, was observed in all but the four most basal genomes. Similarly, we have previously described a single nucleotide insertion that creates a frameshift in the *wbiI* gene of some P314 isolates, which is correlated with a loss of the O-antigen [57] and occurred on branch 6 (S2 Fig).

### Phylogeny

The most parsimonious phylogenetic tree was constructed using 126 core SNPs and has a consistency index of 0.97, indicating character changes that occurred only once, with the exception of two homoplastic SNPs (Fig 3). Consideration of homoplasies and their location in the phylogeny is important for evaluating the degree to which genetic inheritance is clonal or involves lateral gene transfer (LGT) [58]. The high consistency index for this phylogeny and the distant chromosomal location of the two homoplastic SNPs is highly suggestive of a clonal mode of inheritance with no evidence for recombination. This is in stark contrast with the overall *B. pseudomallei* population where diversity, and presumably adaptive evolution, is driven primarily by LGT [19, 59]. This tree is also likely to be a highly accurate reconstruction of the true evolutionary patterns as trees constructed with core genomic SNPs have been shown to be highly evolutionarily stable and yield accurate trees for clonally replicating populations [59–61]. For clonal populations such as this, phylogenetic accuracy is best measured by homoplasy metrics rather than bootstrapping, which tends to underestimate support for nodes supported by a small number of highly stable SNPs [61]. The addition of 36 non-core genome SNPs, 2 single nucleotide deletions, and 15 large deletions as evolutionary characters to the tree increased phylogenetic resolution (Fig 3). Most of these additional genomic characters can be mapped to a single point on the tree; the independence of each large deletion event was inferred by different breakpoints even where deleted regions overlap across strains. The single exception to this is a single nucleotide deletion in two distantly related genomes and must therefore involve two independent events.

We used the genome of K96243 as an outgroup for rooting this dataset. Given the genomic closeness of the environmental isolates, we could not assume that they would form an outgroup as a diverse infecting inoculum could result in environmental strains nested among the clinical isolates. Indeed, diverse inoculating populations of *B. pseudomallei* have been documented [62, 63]. Likewise, the first isolate recovered from the infection (MSHR1043) cannot be assumed to represent an outgroup if the infecting inoculum was diverse, or if *in vivo* evolution had already generated diversity. The rooting of this phylogeny and monophyly of the clinical isolate clade suggests that the infection evolved from a recent single hypothetical ancestor that was likely present in the inoculum. This does not preclude the possibility of a diverse multiple bacteria inoculum where all but one lineage went extinct before the first isolates were collected. In fact, the initial isolate was not the most basal in this phylogeny and suggests that early diversity is not represented in this sample set. The rooting also shows that some of the sampled environmental isolates are closely related to the clinical isolates and differ from the initial infecting strain by only a single SNP. *In vivo* diversity was observed before the collection of the first clinical isolate (MSHR1043), which is not among the most basal isolates in the tree.

Throughout the tree, numerous SNPs and deletions (discussed above) are observed that are specific to lineages that went extinct. In most cases, these lineages only contain single or small groups of isolates, suggesting limited fitness. However, one clade that went extinct contains 13 isolates, suggesting some degree of ephemeral fitness for this subgroup. The large number of isolates sequenced, particularly at the beginning of year 4, allowed us to sample lineages and diversity that would otherwise be concealed. Only two of the many nascent lineages persisted beyond year 4 and continued to diversify. After >16 years, all the extant isolates are derived from these two long-lived lineages (LLL1 and LLL2), and in LLL1, two sublineages persist (Fig 3).

Of the 126 SNPs used to construct the phylogeny, only 24 were synonymous with the remaining 112 changing an amino acid or generating a nonsense mutation. The high nonsynonymous to synonymous ratio (4.7/1) was also observed when only SNPs in LLL-1 (33/7 = 4.7) and in LLL-2 (36/6 = 6.0) were considered. The relatively high number of nonsynonymous SNPs has been seen in other pathogen outbreak populations [64, 65] and has been attributed to either diversifying selection or insufficient time for purifying selection in a young clonal population.

### Mixed genomes

During the course of SNP discovery and evaluation of deleted regions, we identified multiple isolates that showed apparent genotype mixtures. Though isolation streaks were used, the identification of mixtures was not a total surprise as we minimized the number of passages to reduce the chances of additional *in vitro* mutational events. In addition, our level of analyses focused on rigorous identification of even the smallest differences among genomes and the ability to observe low-level read contamination. In two cases, we were able to separate mixtures into minor and major populations based upon SNP frequency in the read data and determine their respective phylogenetic position (Fig 3) by considering which SNP loci showed heterogeneity among sequencing reads. Evidence of strain mixtures was most clear in the deletion matrix (Fig 2), where four instances of mixtures were identified. Even when a dynamic filtering approach was employed to reduce as much contamination noise as possible, intermittent and low-level coverage was still evident in deleted regions for some isolates. Such mixed isolates might have an impact on phenotypic attributes.

### Phenotypic Characteristics

Colony morphology of individual isolates could be described by one of three categories: wild-type, small colony variants (SCVs), and fastidious (S8 Table). As we have previously reported [21], wild type colonies show robust growth and are easily recognized as *B. pseudomallei*. SCVs, however, are difficult to recognize as *B. pseudomallei;* even after 48 h of growth, colonies were typically only <1 mm in diameter. Smaller still were isolates categorized as fastidious; after 72 h, these isolates could barely be discerned on agar plates, appearing as a nearly clear sheen in the primary zone of inoculation with few, if any, pin-prick sized colonies in other areas.

The change from wild type to SCVs arose during the evolutionary history of these isolates with SCVs being present in sputum samples as early as 301 days after the first isolate (MSHR1043) was collected (Fig 3, S8 Table). SCVs have persisted throughout P314’s infection, they occur in both long-lived lineage 1 and 2 (Fig 3), and were isolated from different sample types (i.e. nose and throat swabs, various locations in the lung; S8 Table). Fastidious isolates appear to occur only in long-lived lineage 2 (Fig 3) and are derived from a single sputum sample produced 4,560 days post initial isolate. Interestingly, this sputum sample also produced SCVs in lineage 1 (MSHR7732, 7744, 7734, and 7740) and in lineage 2 (MSHR7729, 7730, and 7735; Fig 3) demonstrating the mixed phenotypic and genotypic nature of this population at this time point. It is unclear if fastidious isolates have developed in lineage 1 as the most recent isolates in this lineage (MSHR9790, 9791, 9509, 9787, 9511, 9788, and 9789) have not been characterized.

Loss of motility was observed in 85 of 110 clinical isolates which include 72 SCVs and all 13 fastidious isolates (S8 Table). All wild type morphology isolates (n=18) were motile. A small subset of SCVs (n=7) that were isolated from P314’s nose and throat between days 395–461 post initial isolate were also motile (S8 Table). Excluding the minor variants of isolates MSHR1360 and 1459 (Fig 3), loss of motility can be mapped to a single branch on the phylogenetic tree that occurs after the phenotypic change to SCVs. This indicates that as late as day 461 there were motile, SCVs still present in the *B. pseudomallei* population (see isolates MSHR1285, 3104, 3105, 1284, 1287, 1290, and 1291; Fig 3). Anecdotally, we noted what appeared to be diminishing motility over evolutionary time in these seven isolates prior to full loss of motility in later isolates. This was indicated by the distance the bacteria appeared to diffuse from the stab line in motility media, with earlier isolates appearing to have a larger diffusion cloud than later isolates. This observation suggests that loss of motility may not be as simple as a single genetic deletion or mutation, which would be implied by motility loss being mapped to a single point on the phylogenetic tree. Instead this suggests a pattern of down-regulation to the mechanism for motility, perhaps interacting with global changes in morphology, which warrants further investigation.

Other phenotypic changes observed in these isolates include the decreased production of the carbohydrates CPS and LPS O-antigen (S8 Table, Fig 3), both of which are important virulence factors [57]. Antibiotic resistance has emerged in many of the clinical isolates and is no doubt selected by the therapeutic regimes used in the futile attempts to cure the infection. One of the most complex phenotypes to change, however, is virulence in animal models.

### Animal studies

To determine if the patient isolates remain virulent in a mammalian host, BALB/c mice were intranasally infected with a select number of *B. pseudomallei* isolates collected from the patient or from the environment at the patient’s home (Fig 6A). The patient isolates were selected to include representatives of the early clades as well as a few later strains, thus enabling us to examine temporal and genetic correlates to virulence changes (Fig 3a). All of the mice infected with the patient isolates survived to 21 days after infection, regardless of the inoculum level. However, all of the mice infected with the environmental isolates succumbed to infection or were euthanized due to clinical score and weight loss by day three after infection. The back-titrated doses of the environmental isolates were between 2600 and 6000 CFU/mouse depending on the isolate. In contrast, the doses of the clinical isolates were higher, between 9250 and 11,040 CFU/mouse, suggesting that the clinical isolates were attenuated even at higher intranasal doses. To better quantify the virulence of the environmental isolates, BALB/c mice were intranasally challenged with a decreasing dose of MSHR1435. All of the mice that were infected with >43 CFU succumbed to infection by day 3 after challenge. In addition, 50% of the mice that received 4 CFU succumbed to infection by 18 days after infection, suggesting that the 21-day LD50 for MSHR1435 is around 4 CFU (Fig 6B). In combination, these data suggest that the clinical isolates are highly attenuated in the BALB/c model even at an infective dose ~1000 fold greater than the estimated LD50 of the related environmental isolate. The phylogenetic position of the tested clinical isolates shows that attenuation evolved rapidly after the infection, as even the earliest clinical isolates had no evidence of virulence in our study.

**Fig 6.**
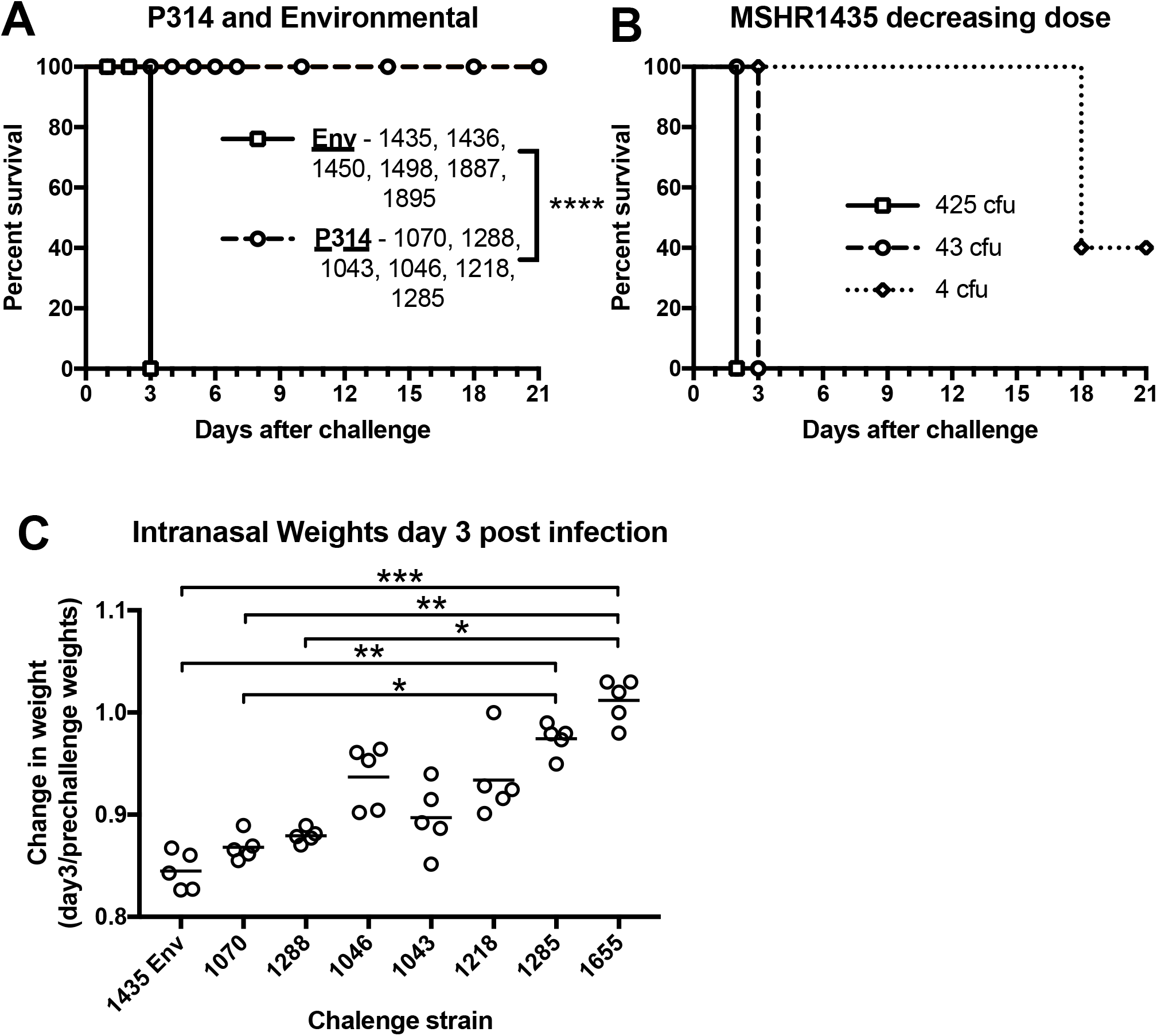
Mortality and morbidity of P314 isolates and P314-associated environmental isolates. A-C) BALB/c mice were intranasally inoculated with *B. pseudomallei* isolates (n=5/group). Mice were monitored for clinical score and were weighed daily for weight loss and if needed were euthanized (A and B). The inoculum for the single dose (A) and the 1435 decreasing dose (B) was back titrated and the CFU/mouse was determined and listed in the figure or below. Back titration results: (A) P314-associated environmental isolates, 1435–2,600 CFU, 1436–3,400 CFU, 1450–2,800 CFU, 1498–3,025 CFU, 1887–6,000 CFU, 1895–6,250 CFU. P314 isolates, 1070–9,250 CFU, 1288–9,550 CFU, 1043–11,040 CFU, 1046–10,350 CFU, 1218–10,150 CFU, 1285–10,450 CFU. Survival significance was determined by Mantel-Cox log rank analysis comparing the individual environmental isolates to the patient isolates. C) Individual percent change in mouse weights at day 2 are shown by dots (n=5/group) and the mean weight is shown by the bar. The isolates are also listed by the relatedness to the environmental isolates cluster (right to left). The change in weight significance was determined with a Kruskal-Wallis test by comparing the change in weight at day 2 for each isolate listed. BALB/c mice were also challenged with 1655 with no mortality but the experiment was terminated early at 14 days (1655) and is not shown. * P<0.05; ** P<0.01; *** P<0.001; **** P<0.0001.

In addition to mortality, we examined morbidity in the mouse model by monitoring weight loss and clinical scores after infection. Infected mice were weighed and scored prior to and after challenge. All infected mice, except those infected with MSHR1655, lost weight until day 2 after challenge. At that point, they either succumbed to infection, had to be euthanized, or started to recover (S3A Fig and S3B Fig). We analyzed the change in weight loss at day 2 to the phylogenetic distance from the environmental isolates (Fig 6C). There is a general decrease in weight loss as the isolates tested become less related to the environmental strain. The average percent weight loss is around 12–15% for MSHR1435 and MSHR1070 then decreases to a 0–2% weight loss for MSHR1285 and MSHR1655. This weight loss is significantly different between these strains (Fig 6). In addition, the weight loss or the lack thereof correlates with an increase or decrease in clinical scores (S2C Fig). In combination, these data clearly demonstrate clinical differences imparted by the environmental versus clinical isolates and additionally suggest that as clinical strains continue to adapt to the host lung microenvironment, mammalian morbidity decreases.

### Virulence gene screen

A set of 551 genes associated with virulence in *B. pseudomallei* [66] were screened across P314 genomes with LS-BSR. Significant variation existed in the conservation of these regions across genomes (a subset of these genes is shown in S9 Table). However, none of these regions were exclusively present in the environmental genomes, suggesting that the differential distribution of these regions fails to explain the virulence phenotype observed in the mouse model. Additionally, no clear trend across isolates used in the animal challenge were observed, suggesting that variation in the conservation of these genes fails to explain differences in animal morbidity.

### Spatial structuring

Removal of the lower left lung lobe on day 1134 afforded the opportunity to collect 38 isolates from various sites and test hypotheses regarding spatial segregation of genotypes. Isolates collected from similar sites are found throughout the tree and multiple clades within the tree contain isolates from multiple, and even distant pulmonary sites (Fig 7). Moreover, isolates from sputum collected before and after the lobectomy are placed in different positions on this tree, suggesting that not only was there a lack of clustering by genotype within the excised tissue, but also that isolates from the excised tissue are representative of the overall lung population. The emergence of the two long-lived lineages with phenotypic and genotypic characteristics raises the possibility of subsequent partitioning of this population. However, at the time of the lobectomy members of these two lineages were present in the excised tissue and had already begun to phylogenetically diverge, showing no evidence of physical separation.

**Fig 7.**
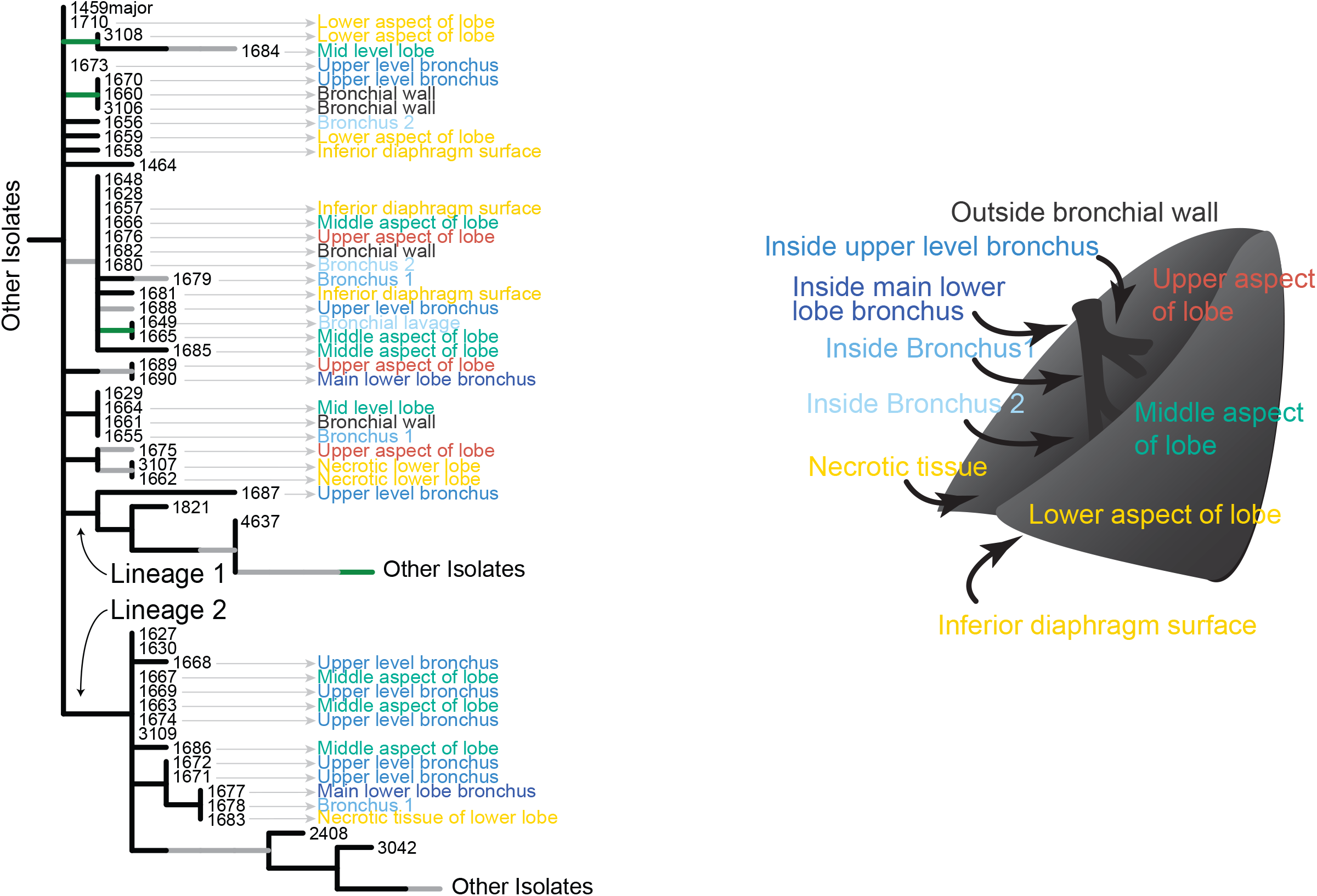
Spatial dynamics. Tree showing location of lobectomy samples suggest one panmictic population. Samples from the same region are found across clades on the tree and multiple clades each contain samples from multiple sites.

### Pan-genome size and diversity

*B. pseudomallei* genomic content is greatly influenced by LGT. However, in the P314 population, there is no evidence for the uptake of exogenous DNA from either conspecifics or allospecifics. The pan-genome accumulation curve (Fig 8) shows a closed pan-genome, whereby the total number of CDSs in the population can be sampled by sequencing only a small number of genomes. Conversely, the number of CDSs shared in all clinical genomes decreases without reaching an asymptotic minimum. The initial increase in the pan-genome size as well as the continued decrease in shared gene content is due solely to the deletion of genomic regions. Consequently, the pan-genome size is limited to the gene content present at the time of infection and the generation of diversity is therefore tied to manipulation of this endogenous gene content. The core genome curve also does not reach an asymptotic minimum, suggesting that continued evolution through novel deletions may be possible for this population.

**Fig 8.**
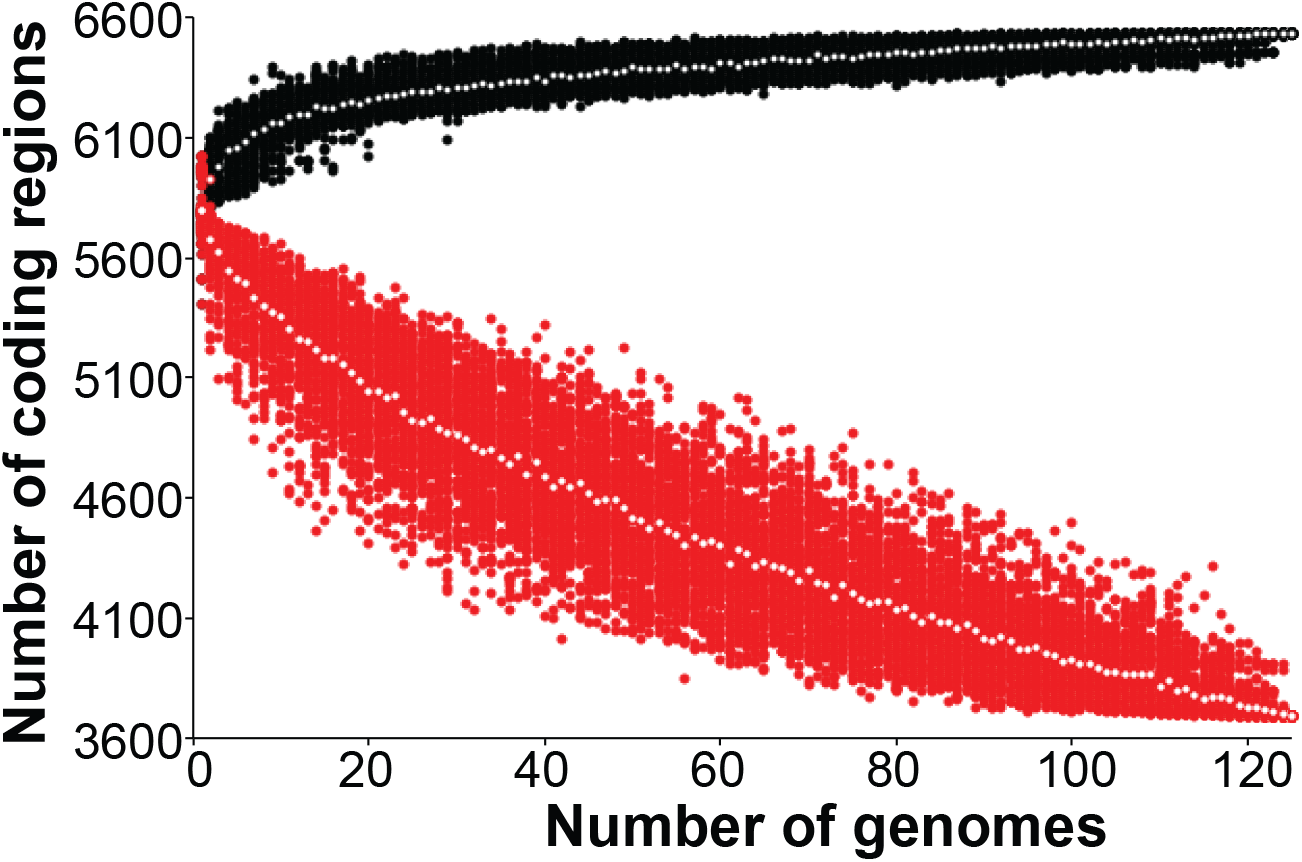
Pan and core genome dynamics. Size of pan (black) and core (red) genome with increased random sampling of genomes show a closed genome and the reduced size of some genomes due to genomic deletions.

### Evolution of diversity

While LGT has not played a role in the evolution of the P314 population thus far, mutation through deletion and point mutation has. While much of the diversity generated within the clinical population is ephemeral, cross sectional diversity increases longitudinally. The phylogeny (Fig 3) shows that even before the first isolate was collected, multiple lineages had diverged through the accumulation of SNPs and deletions. In the first year, 14 isolates were sampled, yielding 9 distinct genotypes (observed OTUs) separated by 14 core and non-core SNPs (Faith’s PD). In the 2nd and 3rd years, the number of isolates collected, the number of observed OTUs, and Faith’s PD for each year remained similar to year 1 values (Fig 9). The population diversity however was not stagnant as the cumulative number of observed OTUs and Faith’s PD increased over this time. In year 4, we collected 47 samples (38 of which were cultured from the lobectomy) but only observed 22 distinct OTUs. Faith’s PD increased to 41 SNPs due to the sampling of multiple short lineages. Through the periods of years 5–12, year 13, and year 17, the number of isolates collected and observed OTUs was also similar to each of the first three years (Fig 9). Although many of the short lineages that were observed in year 4 did not persist, Faith’s PD, however remained high for subsequent periods because of the continued diversification of the two long-lived lineages. The cumulative total of observed OTUs and Faith’s PD increases dramatically at the time of the lobectomy (due to the large number of isolates collected) and continues to increase towards year 17. The difference between Faith’s PD cumulative total and the value observed during year 17 is due to lineages that have gone extinct, although it is certainly possible that some presumably extinct lineages persist but were not sampled.

**Fig 9.**
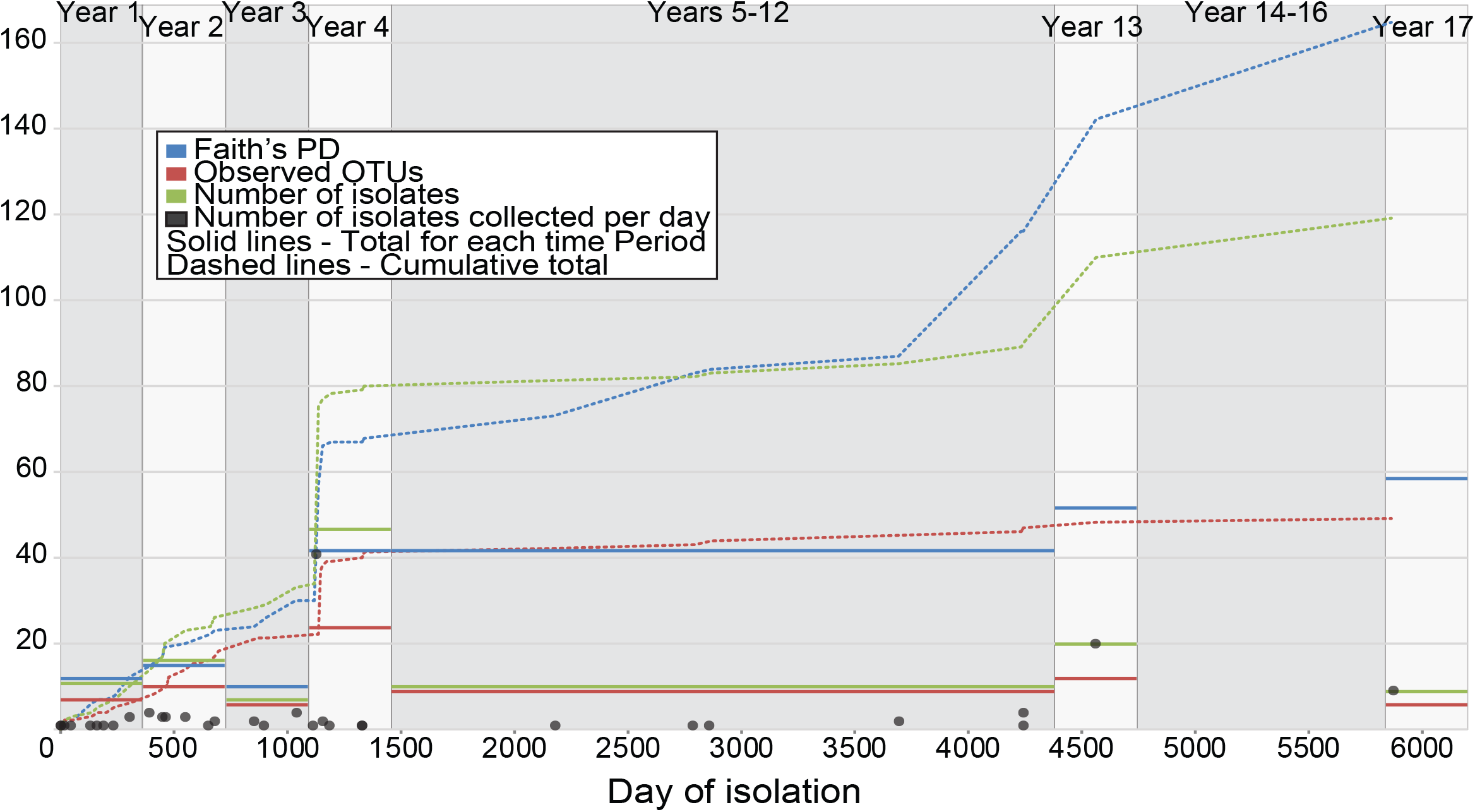
Population diversity dynamics. Estimates of diversity (Faith’s PD and Observed OTUs) in each time period and accumulating over time. This shows that mutations in the first part of year 1 cause a rapid increase in accumulated diversity that continues with time. Continued extinction of earlier genotypes suppresses diversity at each time period. This changes in year 4 when diversity in this time period dramatically increases and is maintained thereafter due to the persistence of two clades. The number of observed OTUs and isolates in each time period remains mostly constant with the exception of year 4 due to the large number of isolates collected from the lobectomy.

### Molecular Clock and Evolutionary Rates

The longitudinal sampling of P314 over a >16-year period and the whole genome data set provide the opportunity to understand the population evolutionary rates in the context of a molecular clock. The median evolutionary rate across the entire genomic dataset was 1.7 × 10^−7^ substitutions/site/year (95% HPD 1.3 × 10^−7^–2.1 × 10^−7^). In agreement with this estimate, it has been previously shown that within patient evolutionary rates of *B. pseudomallei* average 4.9 × 10^−7^ substitutions/site/year [13], *B. dolosa* averages 3.3 × 10^−7^ substitutions/site/year [67], and *B. multivorans* averages 3.6 × 10^−7^ substitutions/site/year [68]. We observed that the effective population size was the smallest at the time of infection, and increased until the partial lung lobectomy occurred in year 4, after which it was stable (Fig 10).

**Fig 10.**
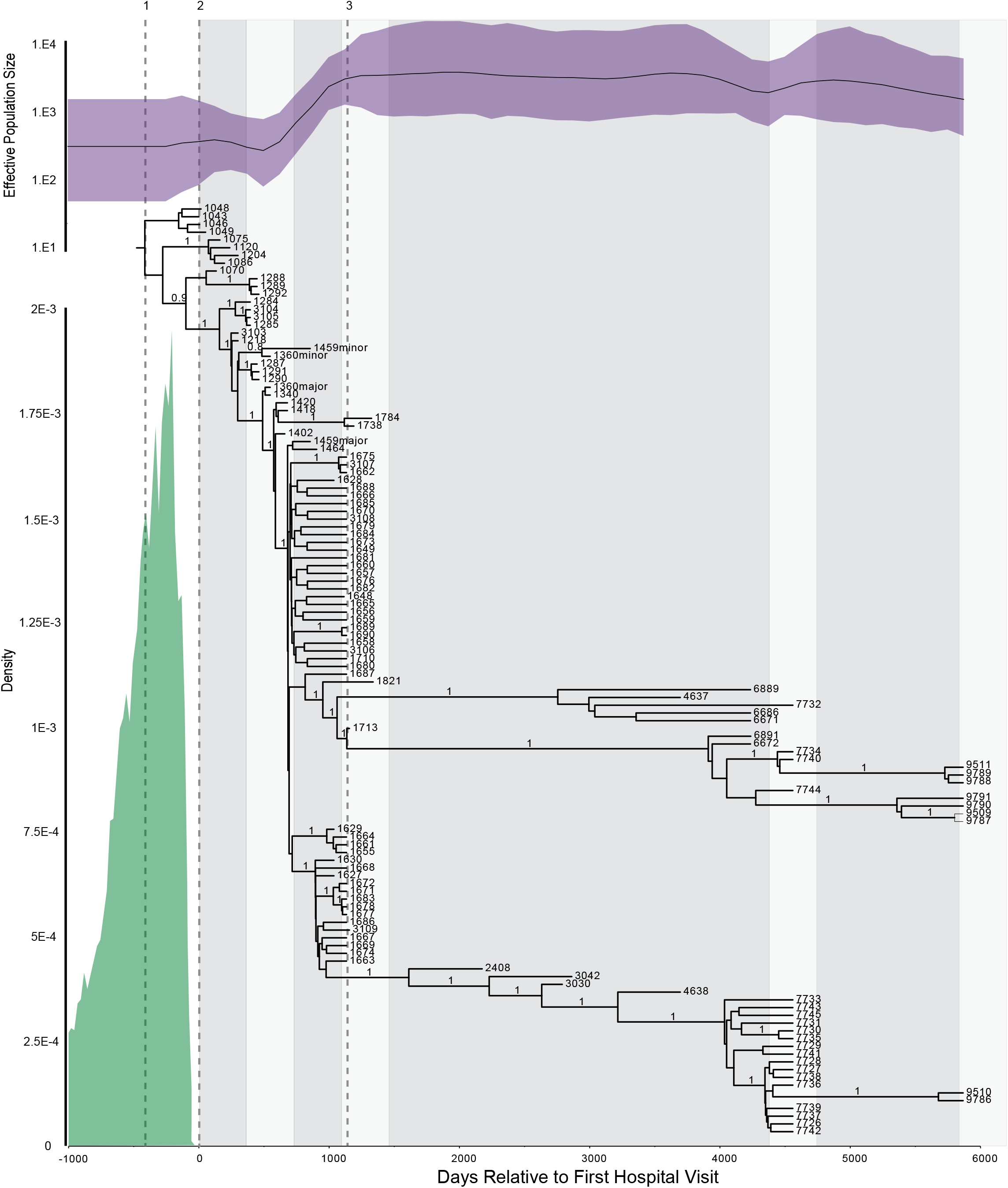
Bayesian estimation of infection date and population dynamics. SNPs from 120 clinical genomes longitudinally sampled over 17 years, were evaluated to determine the estimated time of infection (TMRCA for all clinical isolates) and population dynamics over the course of the infection. The 95% highest posterior density interval is in green and suggests a median time of infection of 343 (mean: 414, 95%HPD: 10–999 days) days prior to the first hospital visit. Posterior probabilities that are greater than 0.80 are indicated for all interior branches. The Bayesian Skygrid (top, purple) demonstrates the fluctuation of the effective population size over the course of the infection. Three important events are indicated: 1) Time of Infection, 2) Time that the first isolate was collected, and 3) Time when lung lobectomy was performed.

The two extant long-lived lineages (LLL) differ in their evolutionary rates. This is visually apparent from a comparison of the branch lengths in Fig 3 where LLL-1 is ~70% of the length of LLL-2. In maximum parsimony analysis, the branch length is determined by the number of character state changes, which in this data set is the number SNPs that occur along each lineage. The SNP evolutionary rates can be estimated from the regression slopes of the branch lengths between the root and each genome against the day of isolation (S4 Fig). The regression slope for the isolates before the LLL bifurcation and the slope for LLL-2 are not significantly different from each other (F=2.244, DFn=1, DFd=96, P=0.1374), consistent with a similar evolutionary rate. In contrast, the slope of LLL-1 is significantly different from the pre-bifurcation slope (F=60.33, DFn=1, DFd=80, P<0.0001) and LLL-2 (F=158.6, DFn=1, DFd=52, P<0.0001). Given these observed similarities and differences in evolutionary rates, we excluded the LLL-1 isolates to calculate the X-intercept (−307.6; 95% CI −240 to −378.2) as an estimation of the time to the most recent common ancestor (MRCA). This represents an approximation of when the patient was infected (days prior to the patient’s first clinical presentation).

We also used a Bayesian approach (Fig 10) to approximate the date of infection. Given the anomalous rate of LLL-1, calculating the time to MRCA with and without these isolates resulted in estimates of median times of 343 (mean: 414, 95%HPD: 10–999 days) and 179 days (mean: 223, 95%HPD 6–558 days), respectively, prior to the patient’s first clinical presentation. The Bayesian TMRCA estimates are earlier than those estimates derived from the linear regression analysis. The regression estimates are based on root-to-tip distances calculated from a maximum parsimony tree, whereas the Bayesian estimates incorporate substitution, site, clock, and demographic models. These dating estimates suggest that this chronic lung infection has persisted for over 16 years.

### Parallel Genome Decay with *B. mallei*

Despite its different taxonomic nomenclature, *B. mallei* is a monophyletic clade within *B. pseudomallei* that has evolved to occupy a specialized niche that is generally associated with chronic respiratory infections (glanders) of equines, though it can also infect other mammals [69]. It has been shown [70] that the evolution of *B. mallei* is strictly clonal and all *B. mallei* are derived from a single ancestral *B. pseudomallei* strain with extensive genome decay that includes large gene deletions. These same characteristics are observed in P314 isolates and indeed, many of the same genomic regions are lost (Fig 5, S6 Table). One particularly interesting observation is that the homolog of the non-ribosomal peptide synthetase, that has undergone multiple deletions in P314, has undergone similar deletions in virulent *B. mallei* genomes (Fig 5).

## Discussion

Bacterial evolution in the natural environment is usually only witnessed through endpoint observations, which greatly limits our understanding of the process (reference). For example, when a pathogen emerges and moves from a reservoir to a susceptible host, a single such ecological jump likely requires a series of sequential adaptive steps but only the resulting pathogen is available for analysis. Endpoint comparisons to understand this process reveal only the mutations that persist in the population in the end state. The nature and extent of ephemeral mutations are lost and the temporal order of many mutations cannot be reconstructed. In this study, we have had the opportunity to sample and characterize a pathogen population in a chronic case of melioidosis across >16 years. We find that the pathogen persisted in the face of therapeutics, evolved, and propagated in a strictly clonal fashion into a commensal that still exists in the human host today.

Chronic infections with *B. pseudomallei* are uncommon but almost always curable and this patient’s original pulmonary disease (non-cystic fibrosis bronchiectasis) is very likely to have contributed to colonization and persistence. Fortuitously and despite high environmental diversity of *B. pseudomallei* in this region, sampling around the patient’s home revealed extremely closely related isolates, providing a surrogate for the bona fide infective inoculum. This reference material allows for precise estimations of the likely ancestral genotype and subsequent *in vivo* evolution. While the patient was initially clinically unwell, the disease pathology progressively lessened to the point that she now has essentially no symptoms that can be attributed to her *B. pseudomallei* infection. Yet she still harbors a *B. pseudomallei* population in her respiratory tract. We have documented sequential genomic and phenotypic changes in the pathogen that are associated with its conversion from pathogen to commensal.

### The progressive adaptation in the respiratory tract niche

Phylogenetic reconstruction of the population’s evolutionary trajectory reveals a highly structured topology where accumulated mutations both persisted and disappeared from the population. The ancestral genotype was rapidly purged from the population and existing genotypes increasingly diverged from the ancestral type. After >16 years, only two lineages (that began to appear in the 4th year), persisted. This is consistent with heavy selective pressure that rapidly purged most variation and resulted in directional evolution. This evolutionary model contains strong evidence of selective sweeps across the entire timeline where single fit genotypes progress in time, with all others going extinct. We do not think that this pattern can be explained through genetic drift exacerbated by small population sizes. This is due to the decadal persistence of the bacterium in a challenging environment, a preponderance of nonsynonymous mutations, and models predicting reasonable population sizes. We believe this demonstrates an evolutionary example involving an active and progressive niche-shift adaptation from a soil-residing pathogen to a respiratory tract environment.

Phenotypic changes are observable in even the earliest clinical isolates, as demonstrated through the comparison of early clinical isolates to environment isolates from the patient’s home. The environmental isolates have nearly identical genomes and are separated in the phylogeny by as few as 2 core genome SNPs and yet have different phenotypic characteristics. While virulence in animals is a complex phenotype, the contrast between the environmental isolates and those from the patient is profound. In the mouse model, the environmental isolates are universally lethal with an exceedingly low LD50 of ~4 CFU (Figs 6A and 6B). In contrast, the clinical isolates all failed to kill at doses >1,000 fold higher (Fig 6A). Our reference to early clinical isolates must be kept in context as we predict that the patient was infected ~2 years prior to clinical presentation and collection of our first isolate (Fig 10). However, from our phylogenetic reconstruction, we can discern the genomic changes and the order in which they occurred in this time period. While diversity was rapidly generated, a deletion in a non-ribosomal peptide synthetase (NRPS) gene was the single change shared by all clinical isolates (Fig 2), indicating that this was the first of all observed mutations. The exact role that this complex enzyme plays is unclear, but it is the sole mutation correlated with early decreased virulence.

Phenotypic changes that could be related to adaptation continued to occur throughout the >16-year timeline. O-antigen is a known virulence factor and a major antigen that triggers intense host responses. We have previously shown that some of these clinical isolates do not produce this antigen and that this is associated with a mutation in the *wbiI* gene [57] (S7 Table). The capsular polysaccharide (CPS) production is greatly reduced in the clinical isolates, though not totally absent (S8 Table). Motility of the isolates is reduced and even eliminated in some clinical isolates, although the genomic basis for this loss is not obvious. In another study (Settles and Keim, unpublished data), loss of motility is associated with the loss of flagellar protein expression. The gross colony morphology changes over time, with some of the most recent clinical isolates having very small colonies and slow growth. And finally, given that morbidity in the mouse model can be estimated by animal weights following infection, we observe that later clinical isolates cause less morbidity than earlier ones (Fig 6C). These examples provide evidence of continuing adaptive evolution even in the later years of this chronic persistence. Some phenotypic changes can be rationalized as avoidance of the host immune response (e.g., O-antigen, CPS, and flagellar antigens). Commensal bacteria are able to co-exist with their hosts in part by not stimulating the inflammatory responses, unlike many pathogens [71]. However, we have little understanding of this niche and its complexity, which limits our ability to understand the full extent and repertoire of selective pressures driving this adaptation.

### The Maintenance of Two Long-Lived Lineages

While selective sweeps and lineage extinction are very common in this population, in year 4, two lineages emerged and persisted for at least 12 years (Figs 3 and 10). Even within these two lineages, selective sweeps and extinction continued to occur, consistent with continued stepwise adaptation. These two lineages however persisted. Relatively large genomic differences exist between these lineages, including differential genome deletion patterns: long-lived lineage 1 is missing region 6d, while long-lived lineage 2 has lost regions 2b and 7 (Fig 5A, Fig 2). Phenotypic differences are also apparent, with long-lived lineage 2 exhibiting very small colony morphology and slower growth. Given these phenotypic and genotypic differences, how and why do members of both lineages persist? Perhaps they are adapting to two distinct niches. The existence of different niches in the respiratory tract would imply differential selective pressures and if subpopulations become partitioned in these niches, could result in spatially adapted genotypes. While limited, the data from year 4 does not support this allopatric model; isolates from extensive pulmonary sampling associated with the lobectomy showed evidence of genotype mixing across pulmonary space and no observable spatial differentiation at a macro-scale (Fig 6). However, massive lineage extinction occurred following the lobectomy, with only these two long-lived lineages persisting. The two evolutionary paths to commensalism mostly occurred in the following 12 years and could involve spatial differentiation in the respiratory tract. With functional mucociliary clearance mechanisms and aspiration, bacterial movement within the respiratory tract is likely, thus differential niche adaptation can occur without spatial segregation.

An alternative to the two-niche spatial hypothesis is of a single niche with differential selective pressures. For example, temporal shifts in host/microbe interactions may vary selective pressures in the same space, allowing for the maintenance of different lineages. Frequency dependent selection due to different host interactions and avoidance abilities can also result in balancing selection to maintain diversity in sympatry. In contrast to competition between lineages, cooperation may also result in balancing selection if both lineages possess different sets of competitive advantages that can be mutually exploited. For example, in nutrient poor environments, differential production of extracellular small metabolites such as siderophores can provide nutrients to conspecifics that would not otherwise be able to obtain them. In a clonally reproducing population, the maintenance of both genotypes could be critical to survival as selective sweeps and background selection will frequently purge alleles (including beneficial ones). Because we have isolates from each lineage, it should be possible to test hypotheses concerning adaptation to particular environmental variables (e.g., oxygen levels).

### Host Adapted *B. pseudomallei* is Evolving Convergently with *B. mallei*

Similar to this population, *B. mallei* is a strictly clonal pathogen derived from a *B. pseudomallei* ancestor. It is also host restricted (found primarily in equines) and genome decay is thought to have been a major evolutionary driver for host-specific adaptation after its derivation from *B. pseudomallei* [70]. In addition to these striking general similarities, even similar genomic deletions are shared by the P314 isolates and *B. mallei* (Fig 5). *B. mallei* is also manifested by chronic respiratory infections but is highly virulent and passed from animal to animal with only a transitory environmental phase. Because *B. mallei* is a clonal derivation from *B. pseudomallei*, it is postulated that an equine host contracted melioidosis, which then evolved into a chronic respiratory disease. Equine pulmonary melioidosis may have occurred many times, but only once did it result in all of the extant *B. mallei* isolates. While both populations (*B. mallei* and P314) adapted to a chronic respiratory tract niche, it is most parsimonious to infer that virulence in *B. mallei* was maintained rather than lost and regained. Likewise, for our chronic patient’s isolates, we can only speculate on the likelihood of continued evolution, perhaps through recombination, that may occur if some members of this population were introduced into the environment (via expectoration). Such convergent evolutionary results can be due to different mechanisms and this human chronic infection provides a single rare example of possible evolutionary pathways towards the emergence of potentially deadly pathogens such as *B. mallei*

## Supplemental Files

**S1 Fig.**
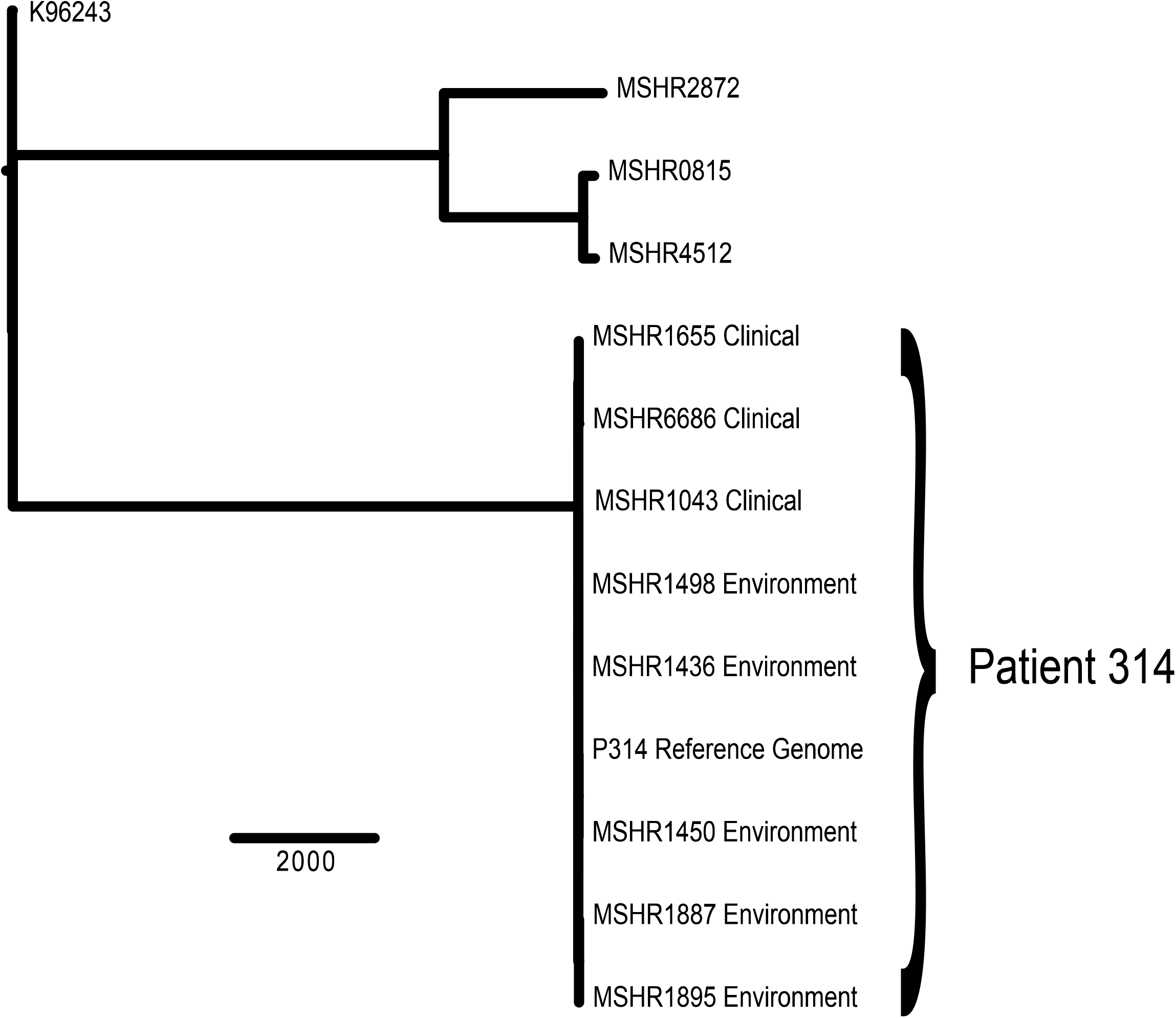
Core genome relationships among ST131 isolates. SNPs (17,359) from core genomic regions were identified and then used to construct a maximum parsimony tree of 11 ST131 isolates’ genetic relationships. K96243 was used as an outgroup for rooting. The two strains (MSHR1043 and MSHR6686) isolated from patient sputum clustered tightly with the six environmental isolates from the patient’s residence. The P314 reference genome is the closed genome of the environmental MSHR1435. The CI of this phylogenetic reconstruction was 0.96, which is indicative of a highly clonal mode of genetic inheritance. Over 16,000 SNPs separate the P314 cluster (clinical and associated environmental isolates) from the next nearest ST131 strains.

**S2 Fig:**
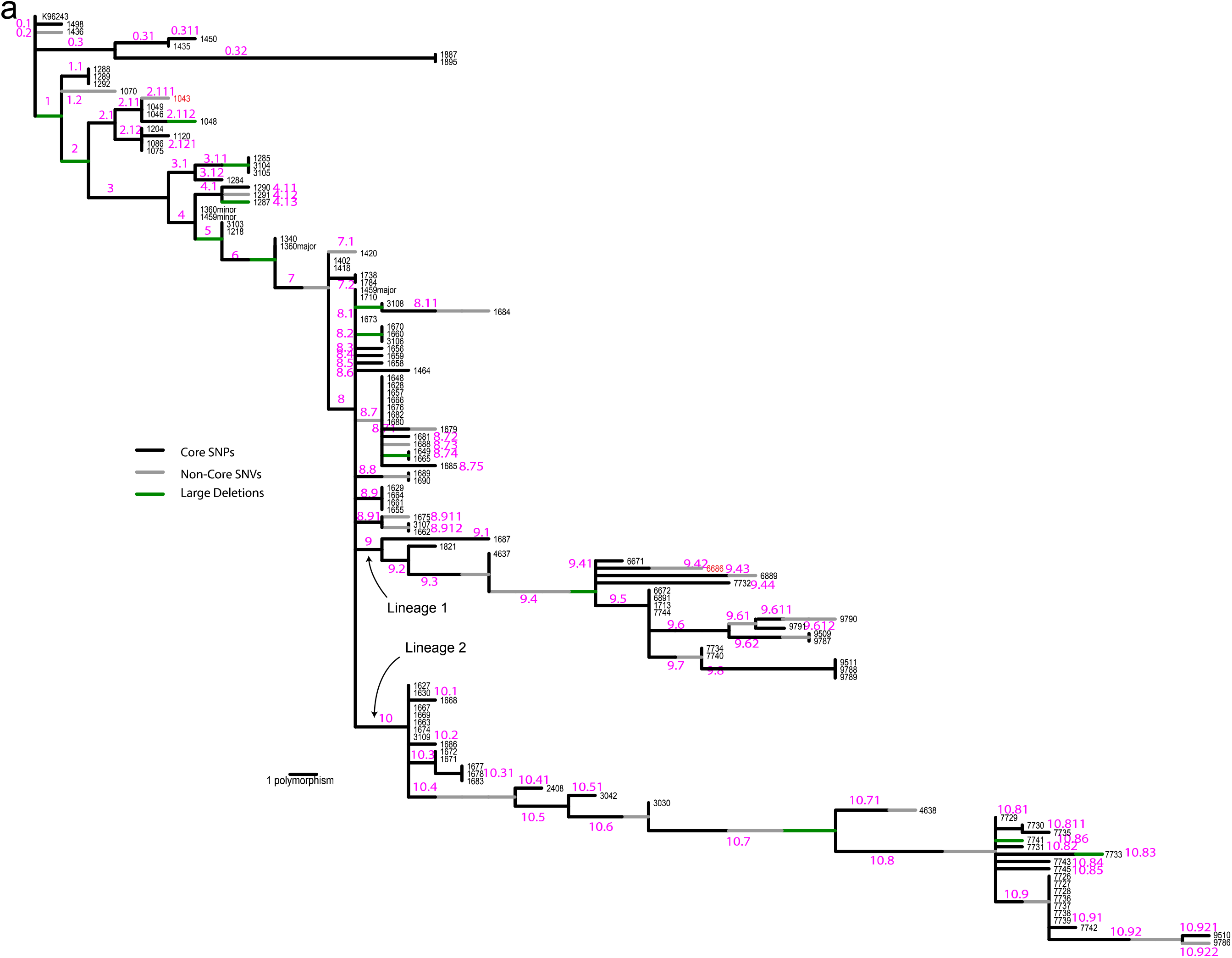
Tree for branch labels. Maximum parsimony tree from Fig 3 with branch labels. Table 1 lists all polymorphisms and includes information on the branch where the mutation occurred.

**S3 Fig.**
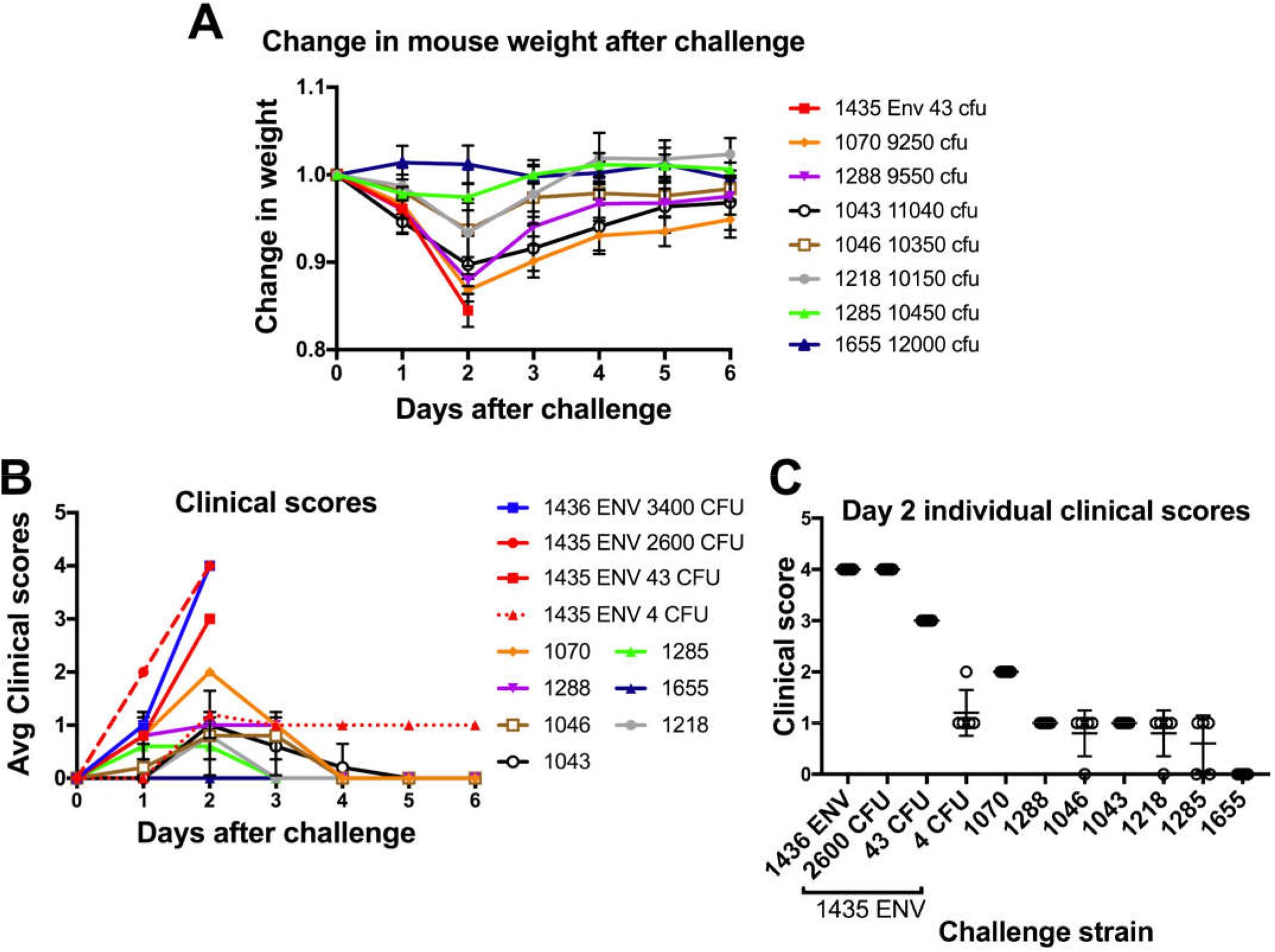
Morbidity of P314 isolates and P314-associated environmental isolates. A-D) BALB/c mice were intranasally inoculated with *B. pseudomallei* isolates (n=5/group). Mice were weighed and clinically scored daily. A) The average change ± standard deviation in weight of the group over time is shown for the initial 6 day period. B) The average clinical score ± standard deviation of the group over time is shown for the initial 6 day period. An increase in clinical score indicate an accumulation of clinical symptoms or an increase in morbidity. C) Individual clinical scores at day 2 are shown by dots (n=5/group) and the mean score is shown by the bar. The isolates are listed by the relatedness to the environmental isolate cluster (right to left).

**S4 Fig:**
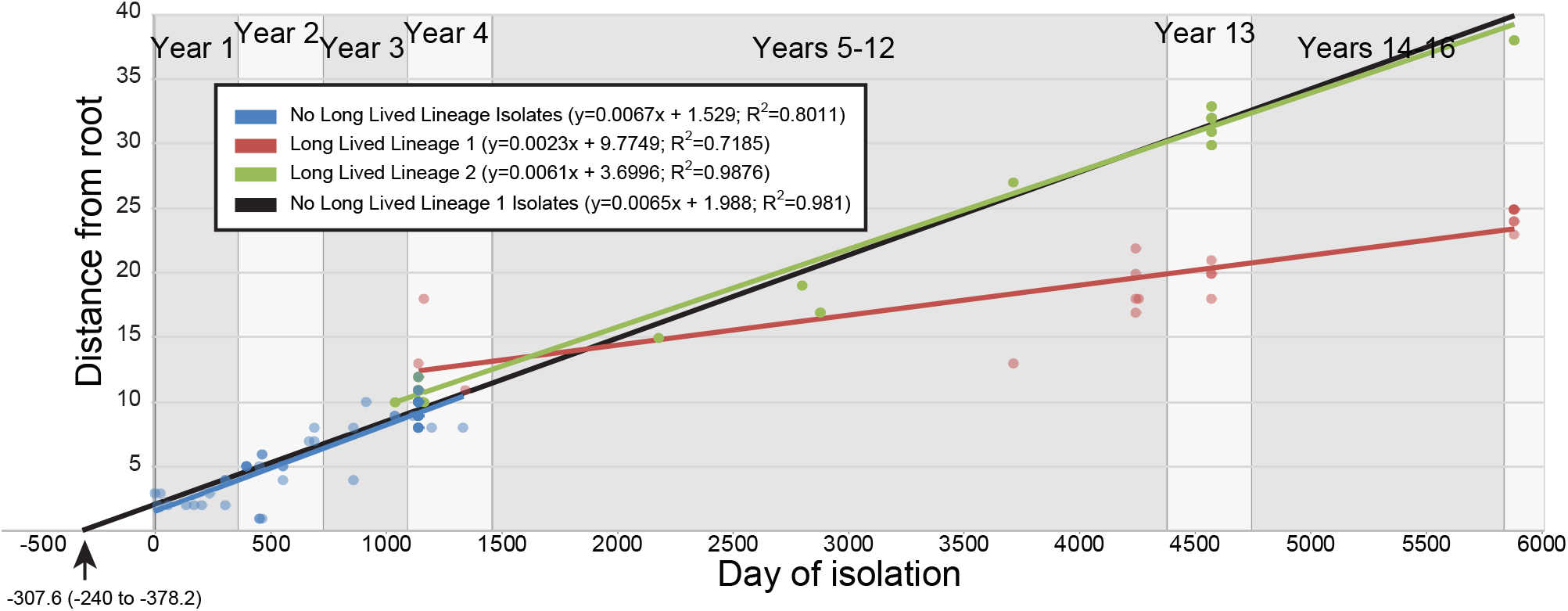
Distances from the phylogenetic root to each genome. Similar slopes for isolates phylogenetically located before the divergence of the two long-lived lineages and those of the long-lived lineage 2 suggests similar rates of evolution. A smaller slope for isolates from the long-lived lineage 1 suggests a lower evolutionary rate, however relatively large root-to-tip distances for the earliest long-lived lineage 1 isolates may indicate a temporary jump in evolutionary rate. Extrapolation of root to tip distances (excluding the long lived-lineage 1 isolates) suggests divergence from a common environmental isolate ~308 days before presenting at the hospital. This is a slightly longer estimate than provided by BEAST (Fig 9) which also utilizes invariant sites to estimate mutation rates.

**S1 Table:**
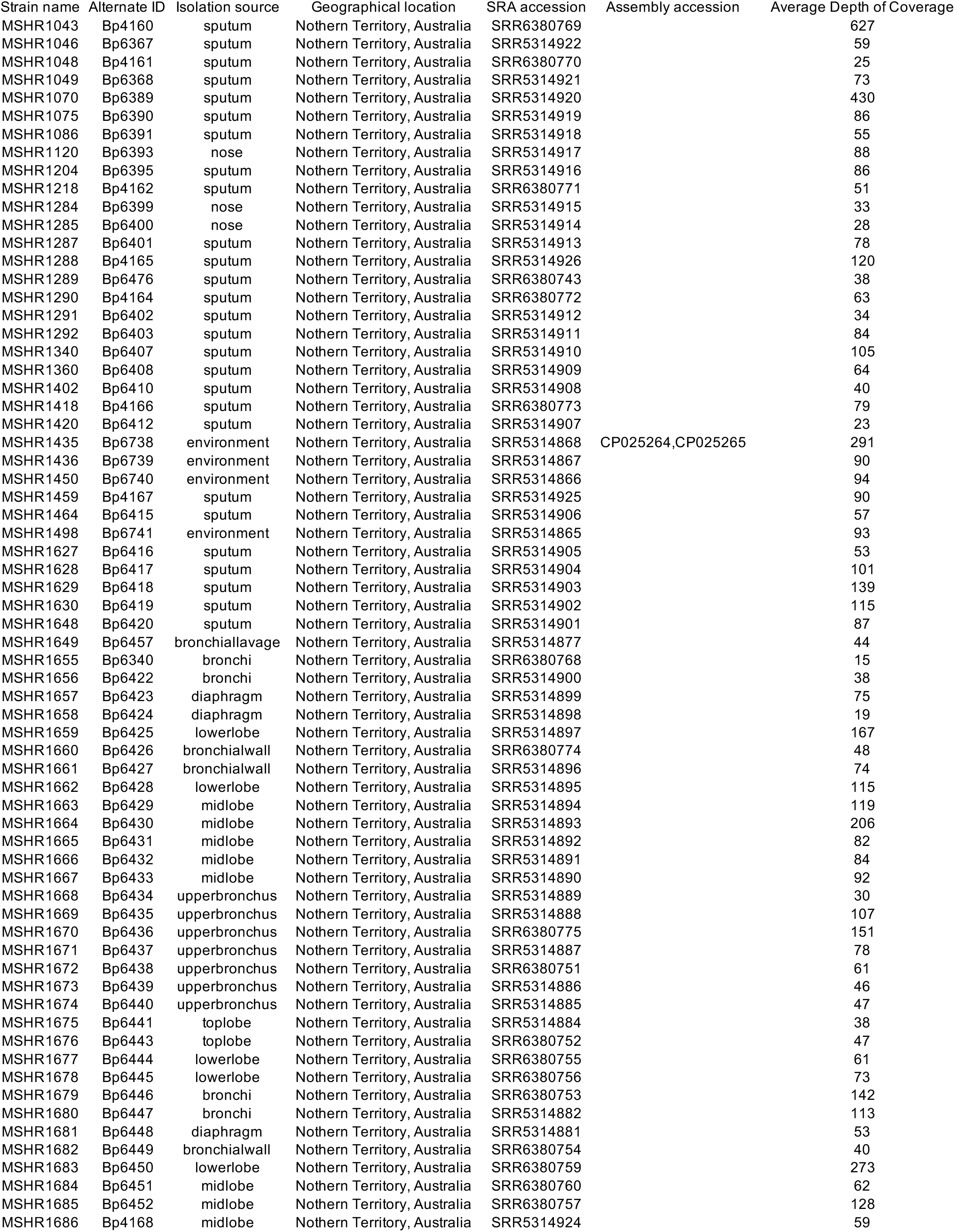
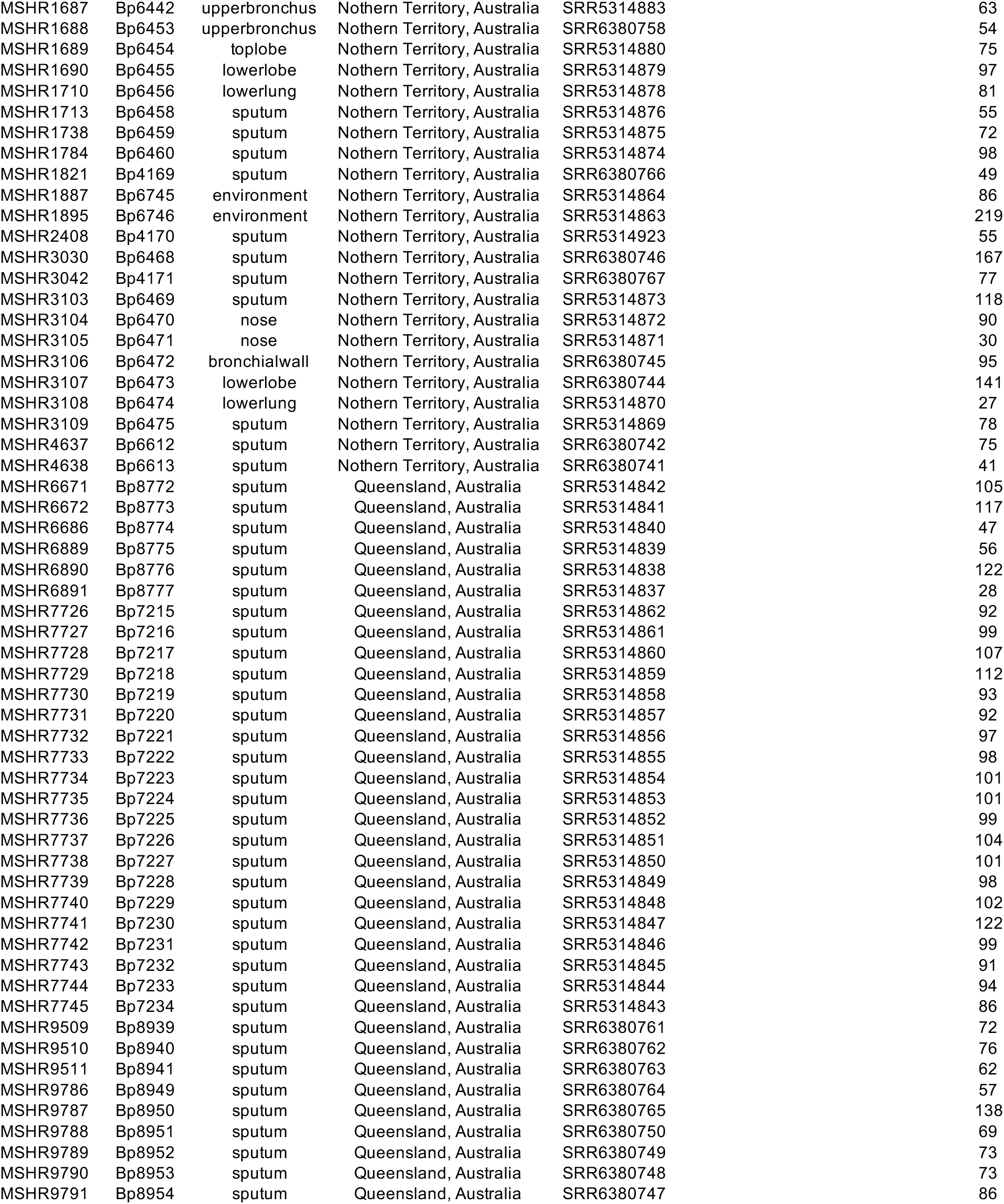
Genome details.

**S2 Table:**
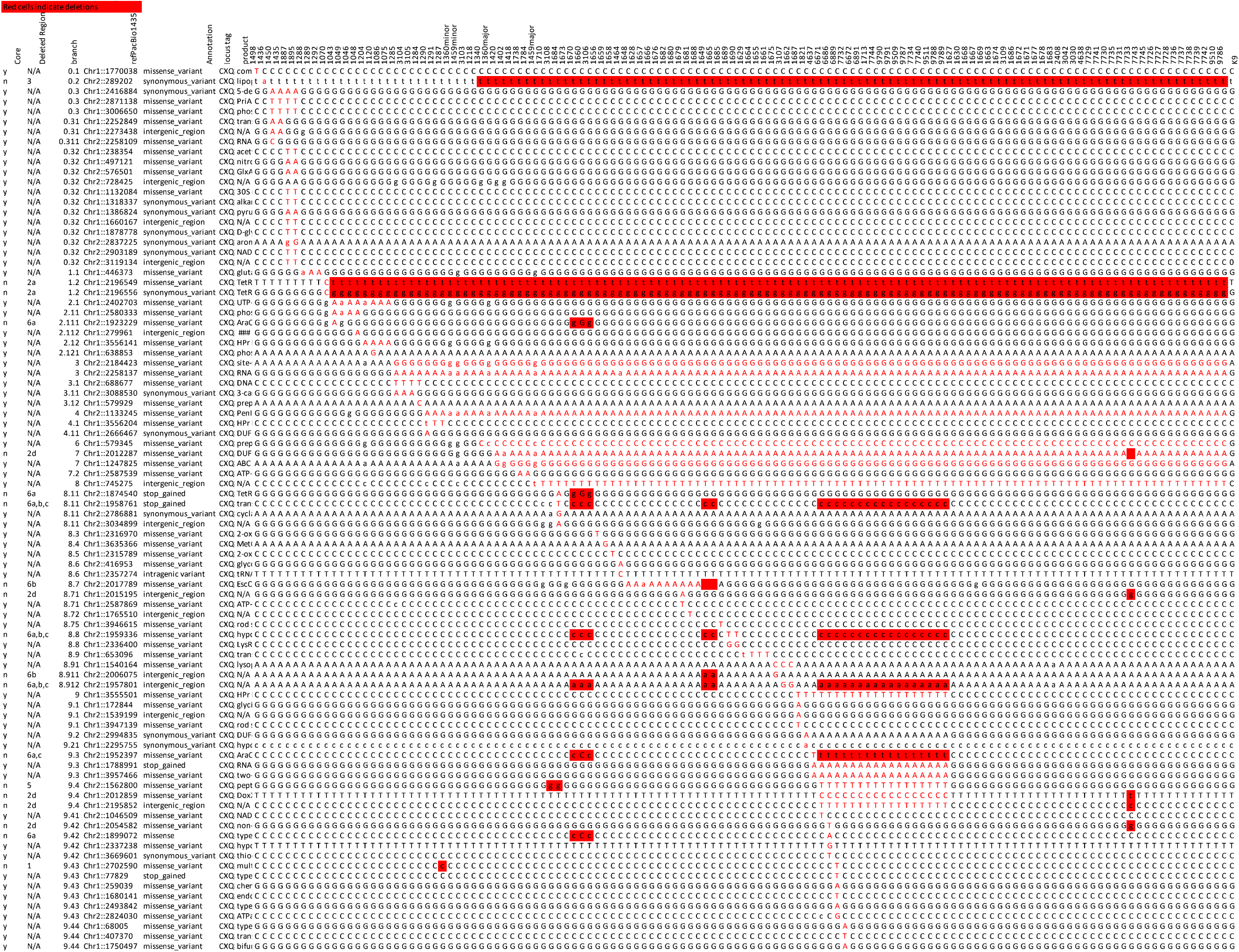
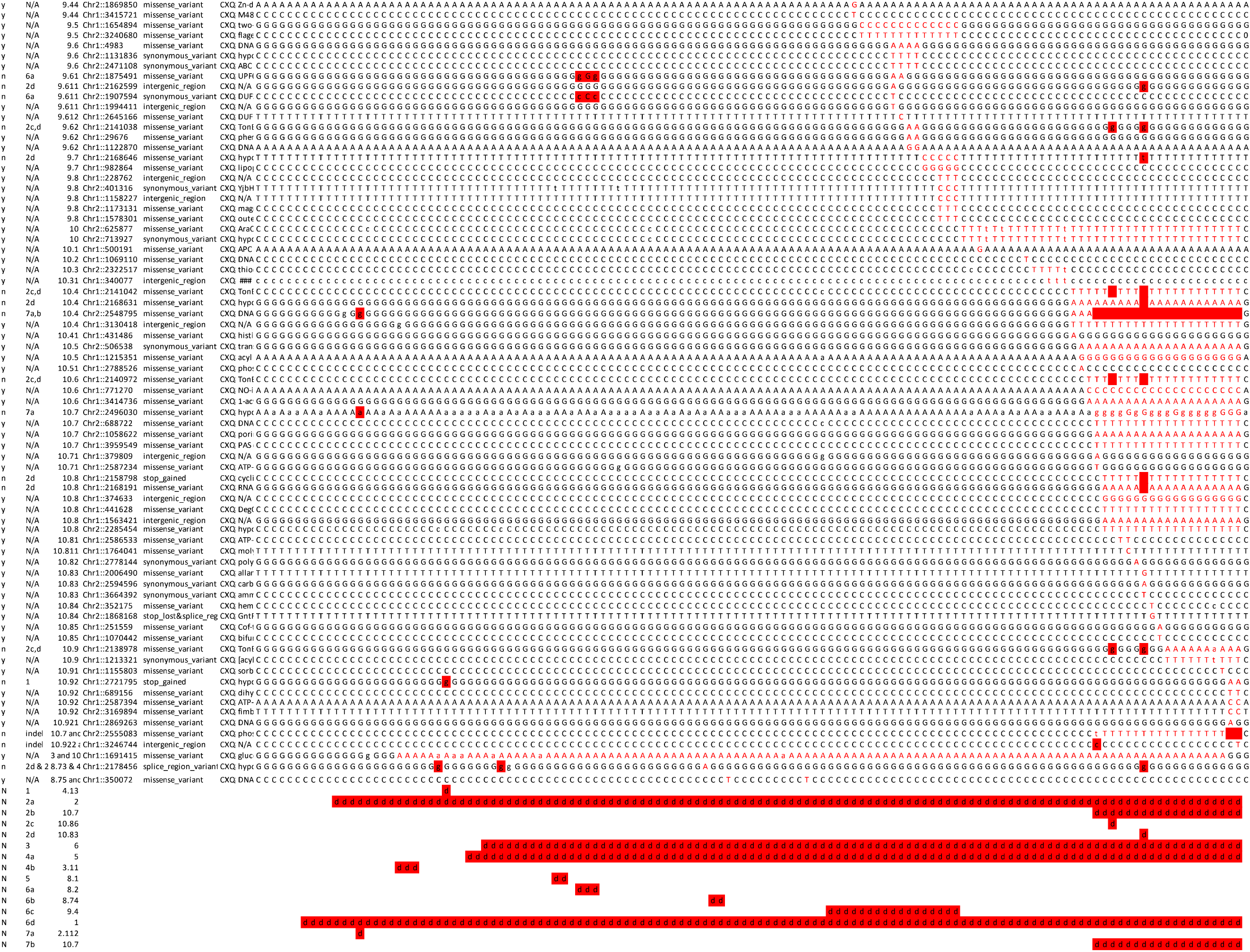
Phylogeny matrix.

**S3 Table:**
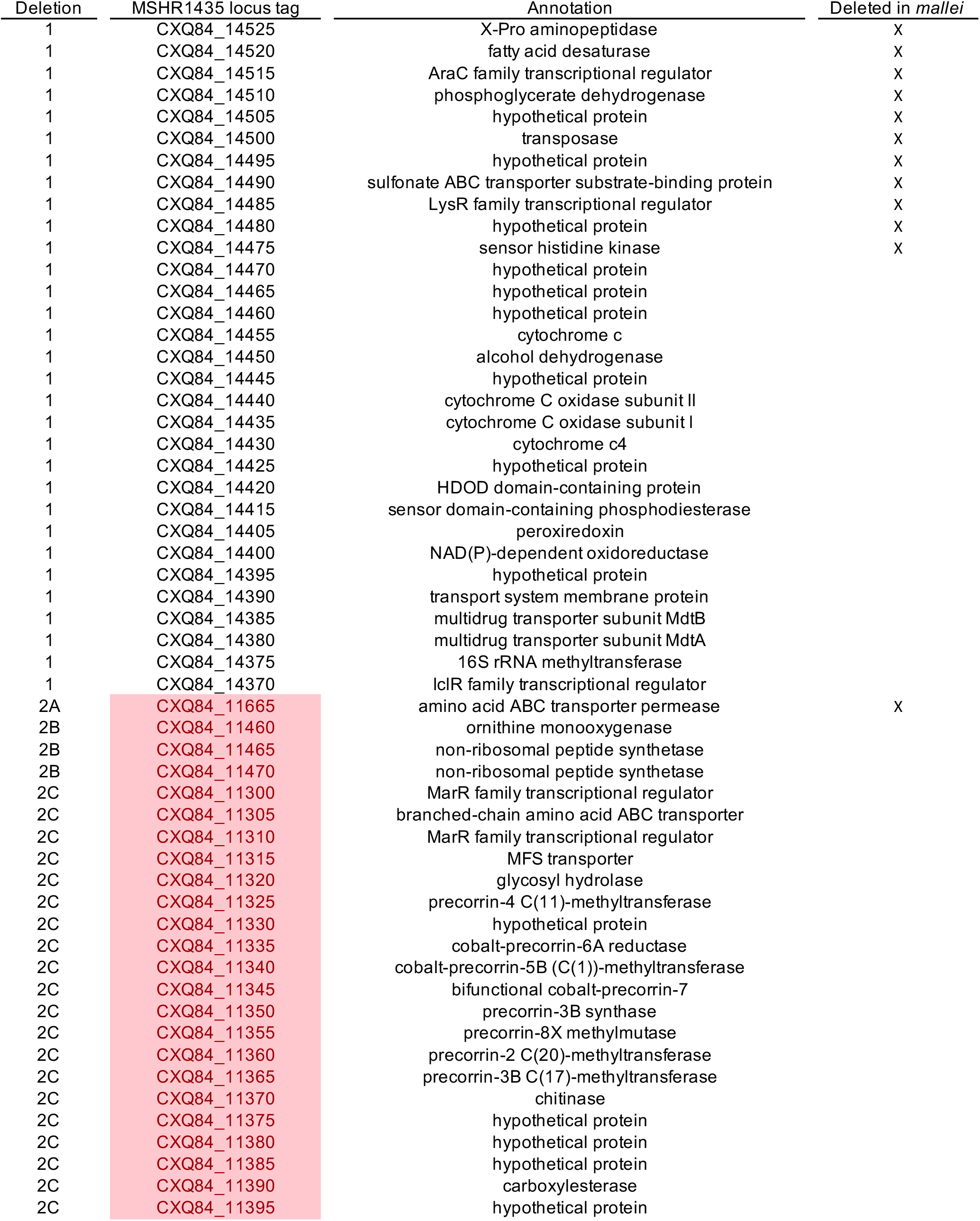
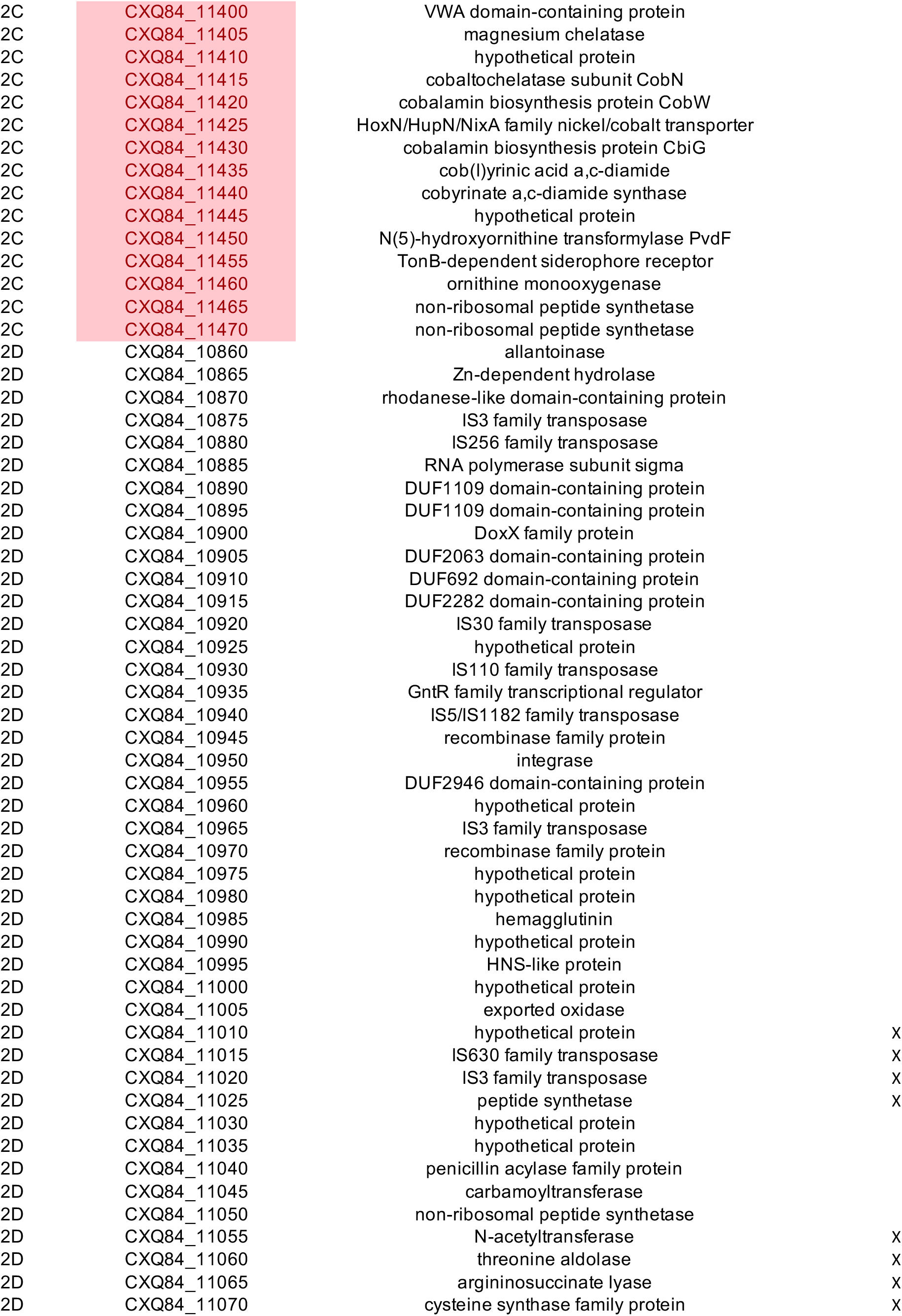
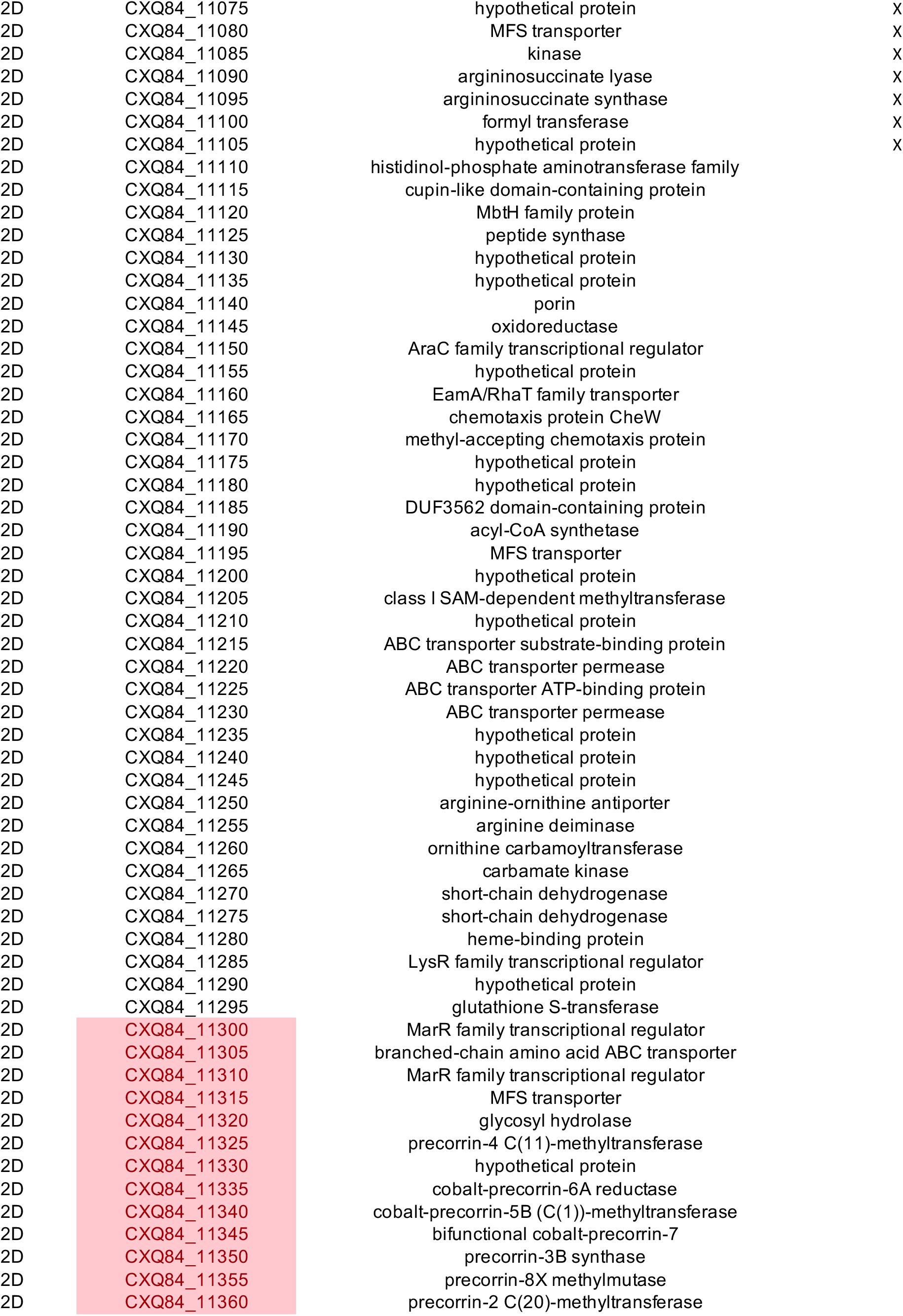
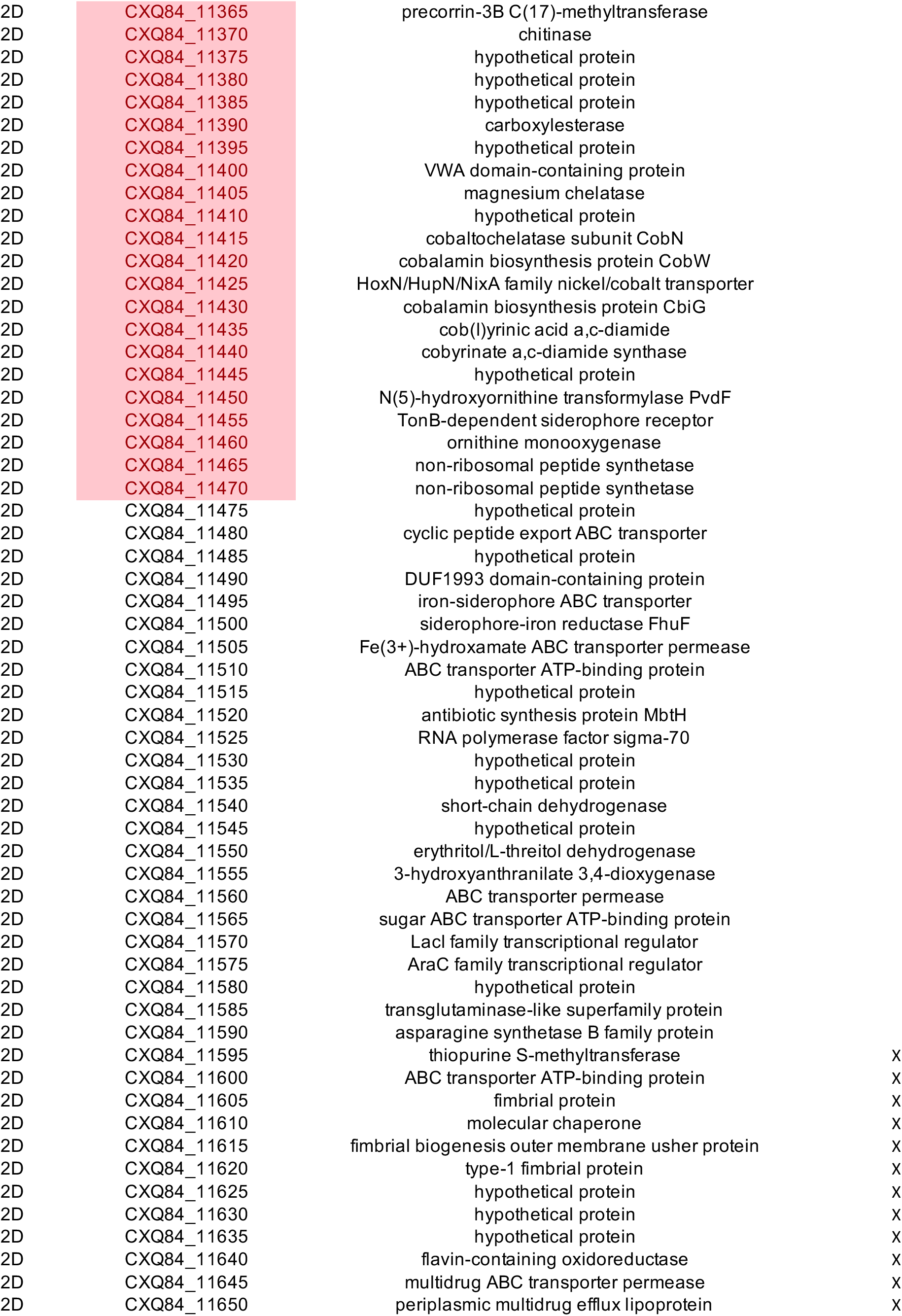
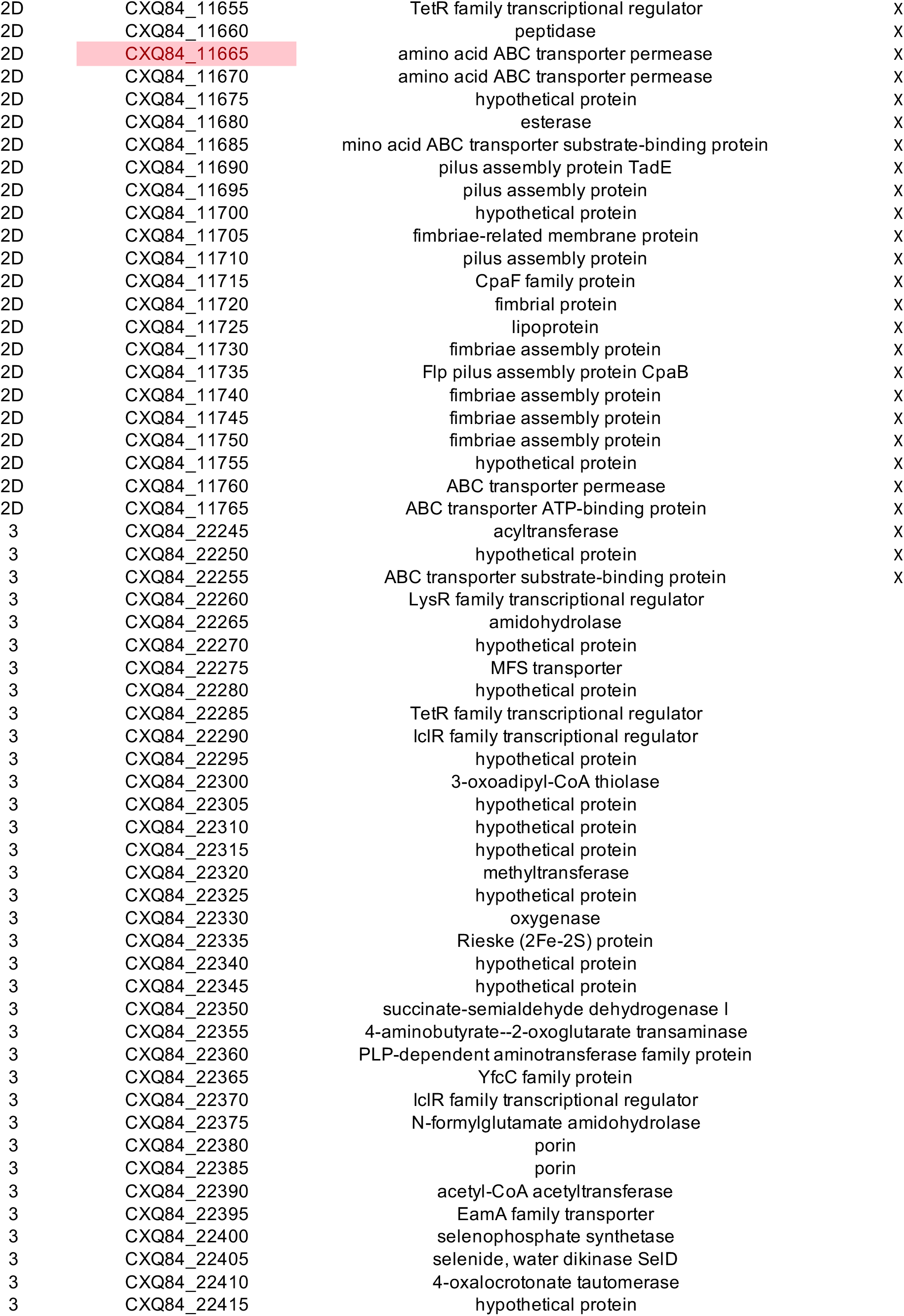
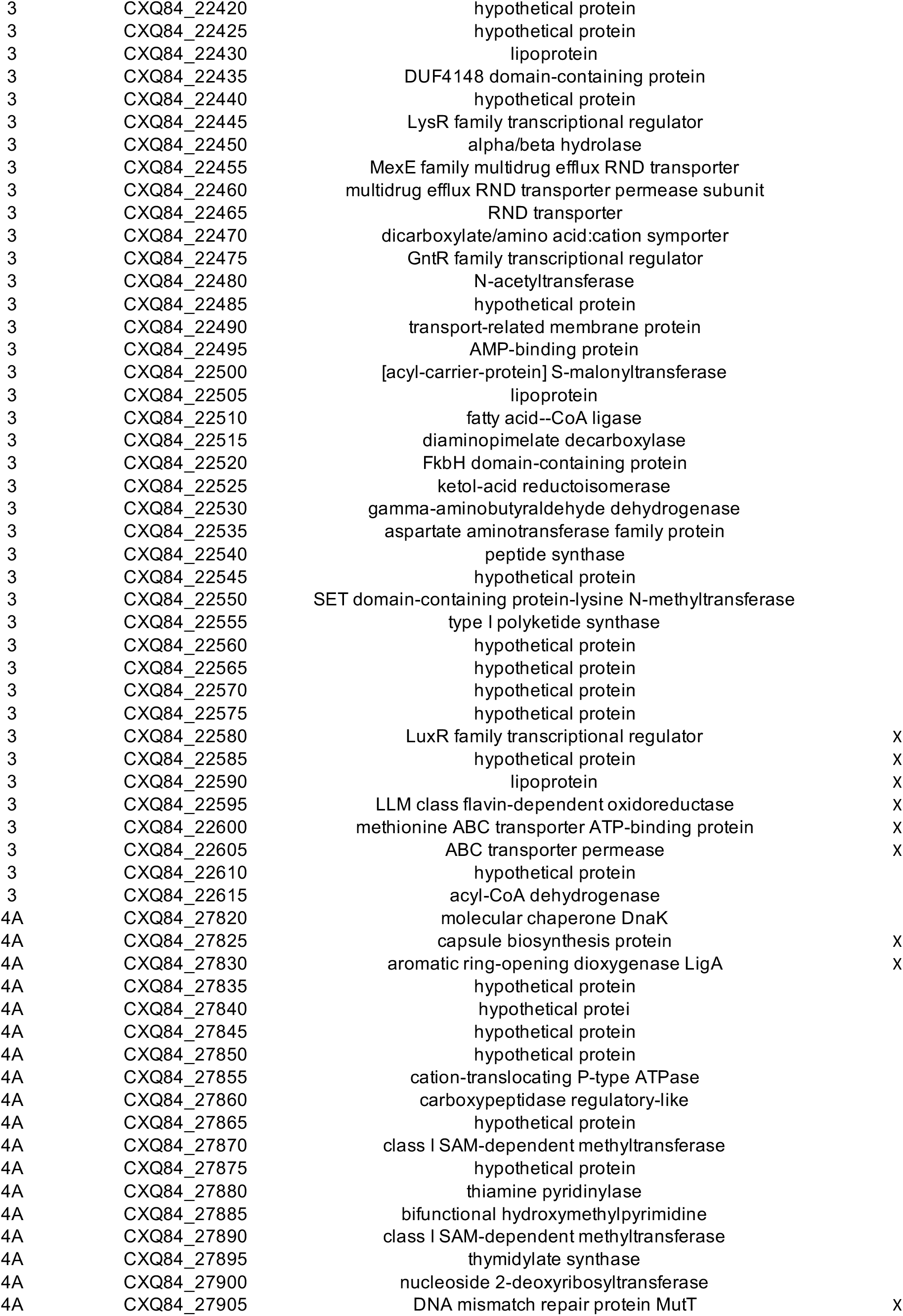
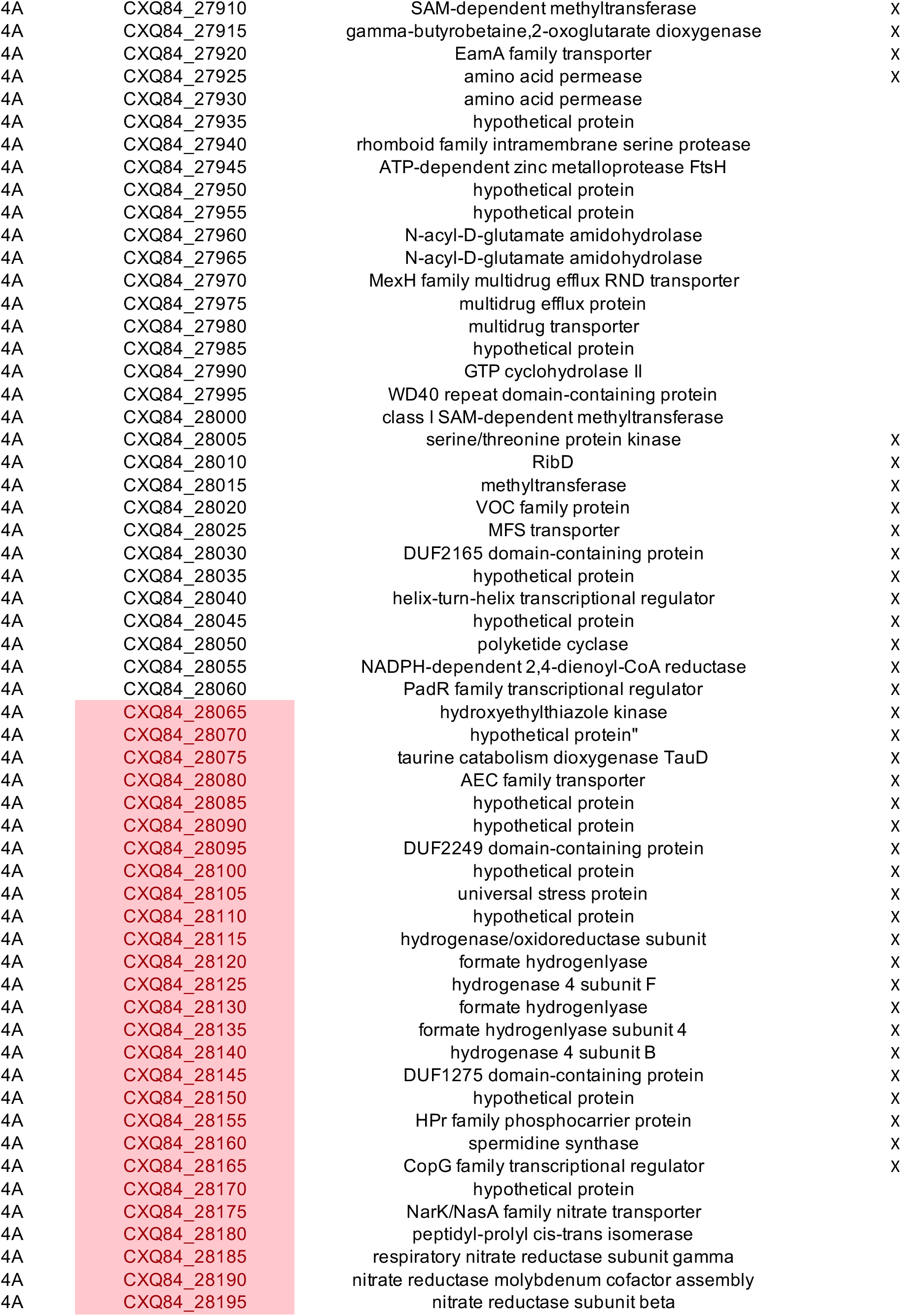
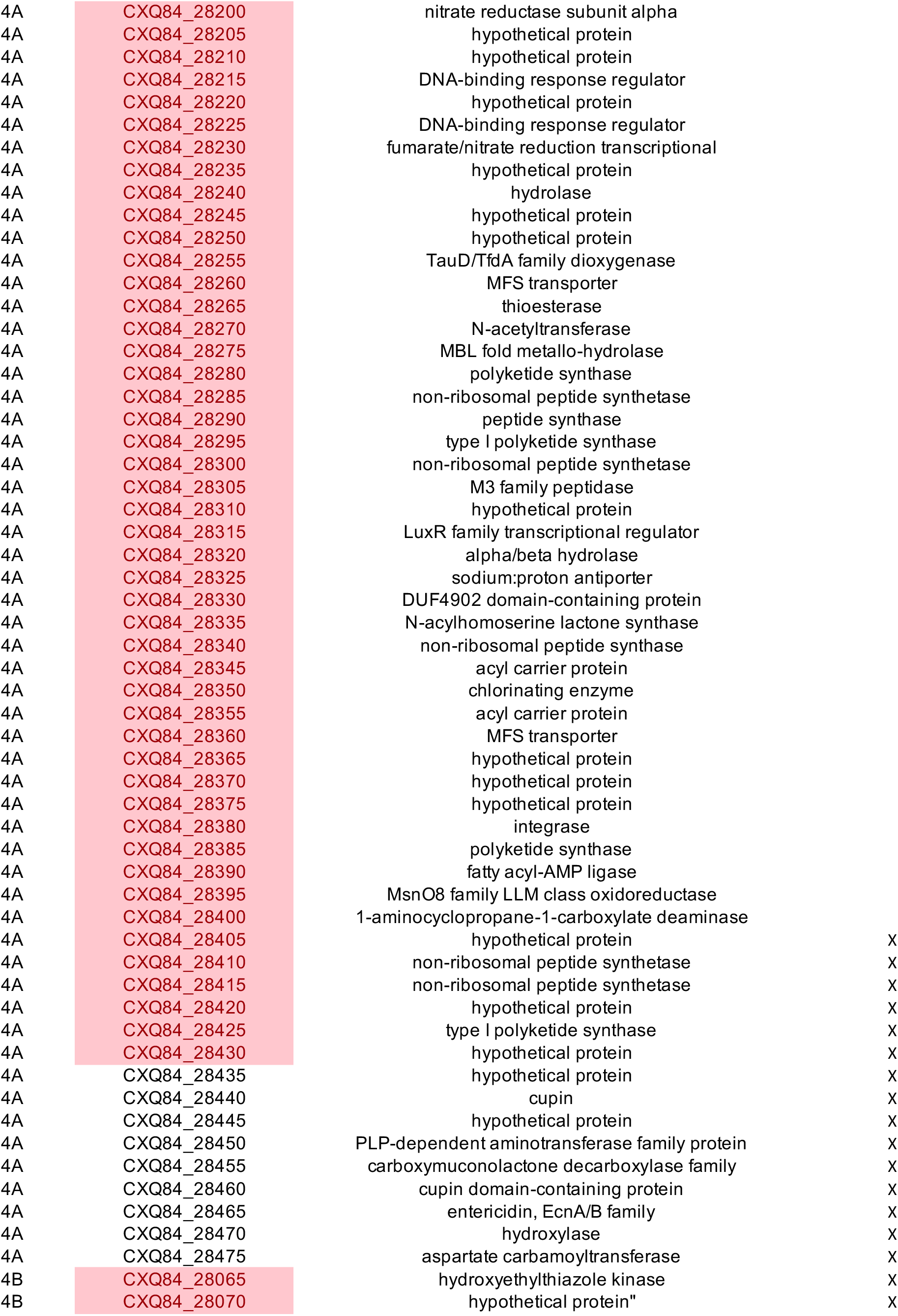
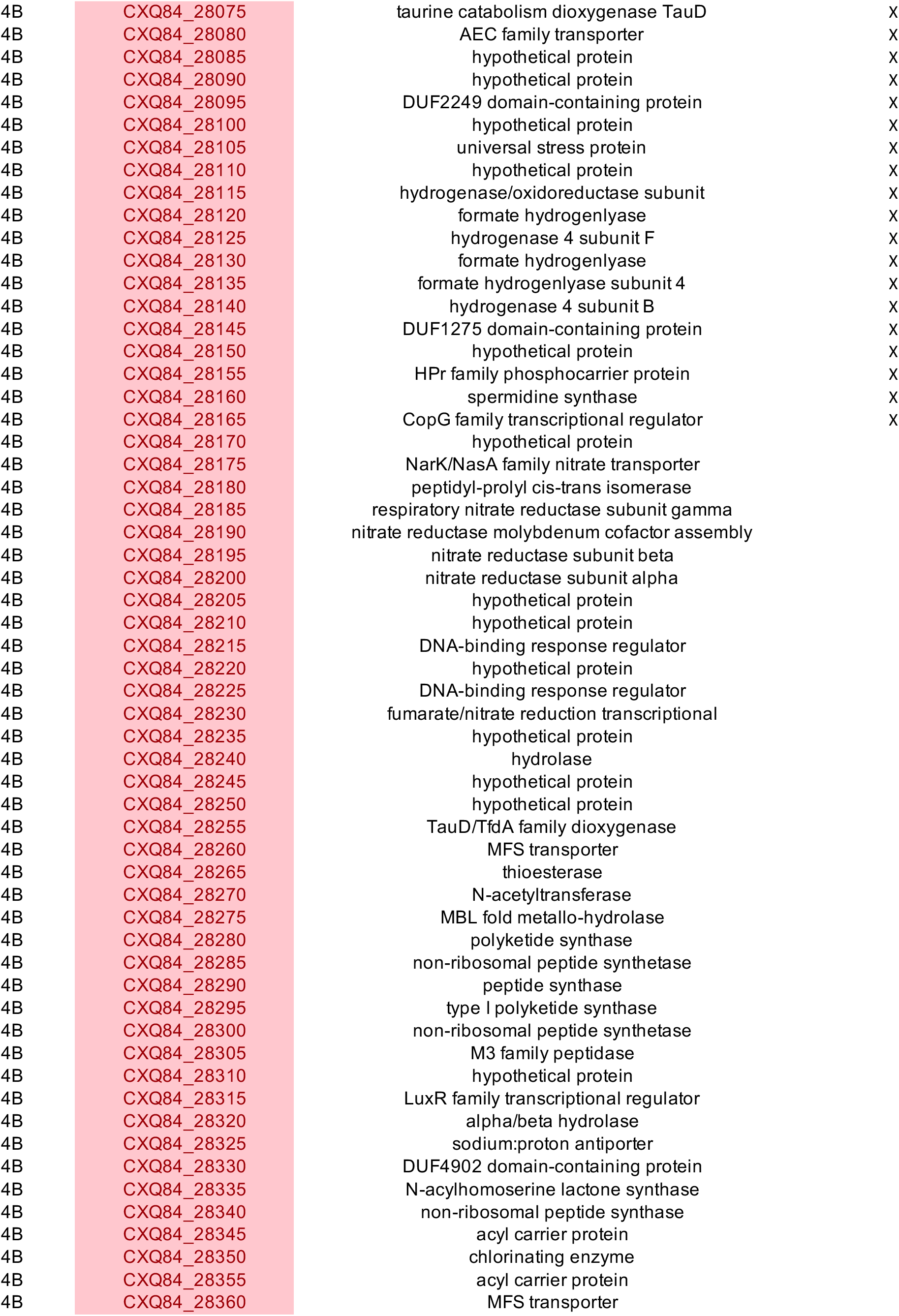
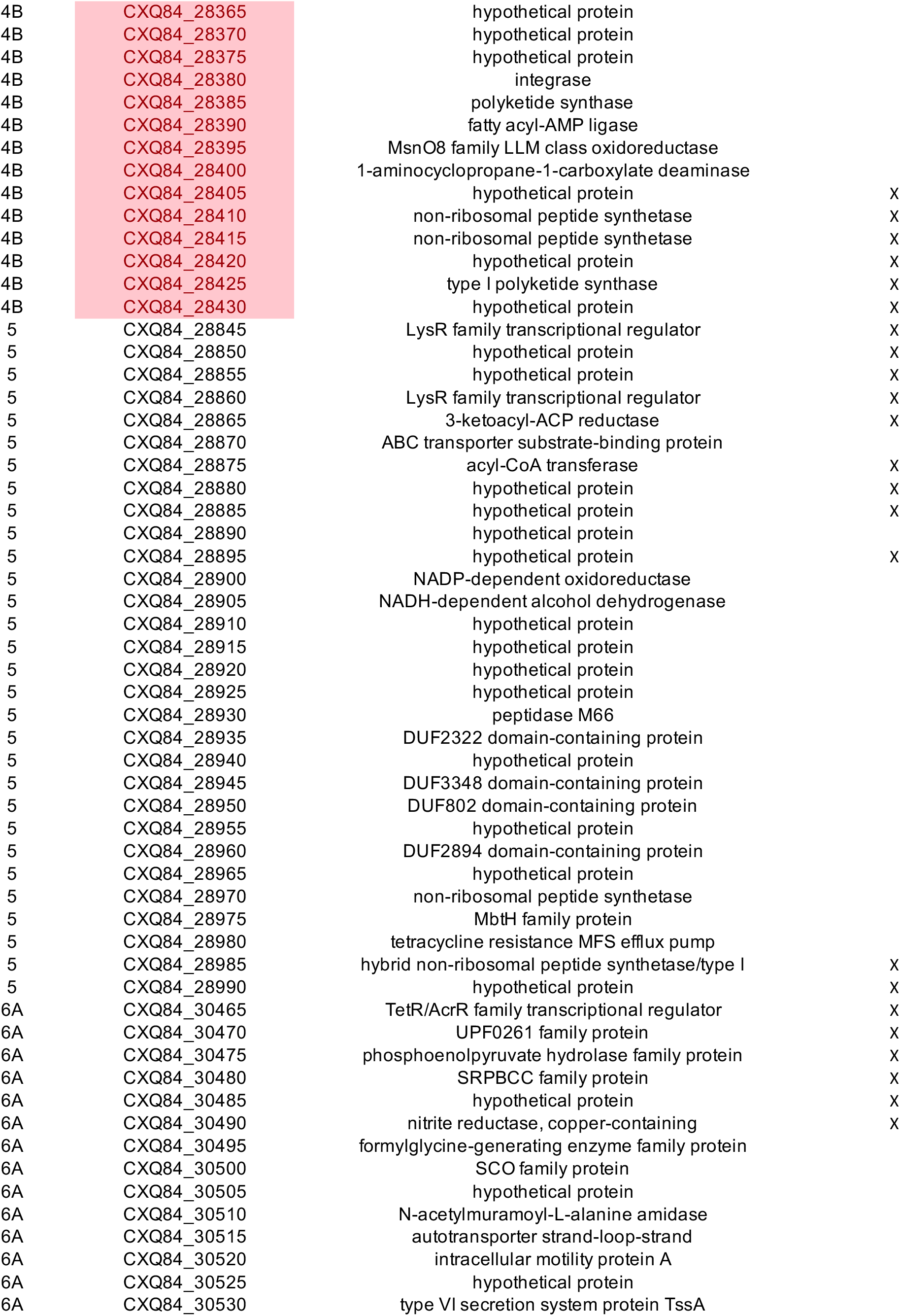
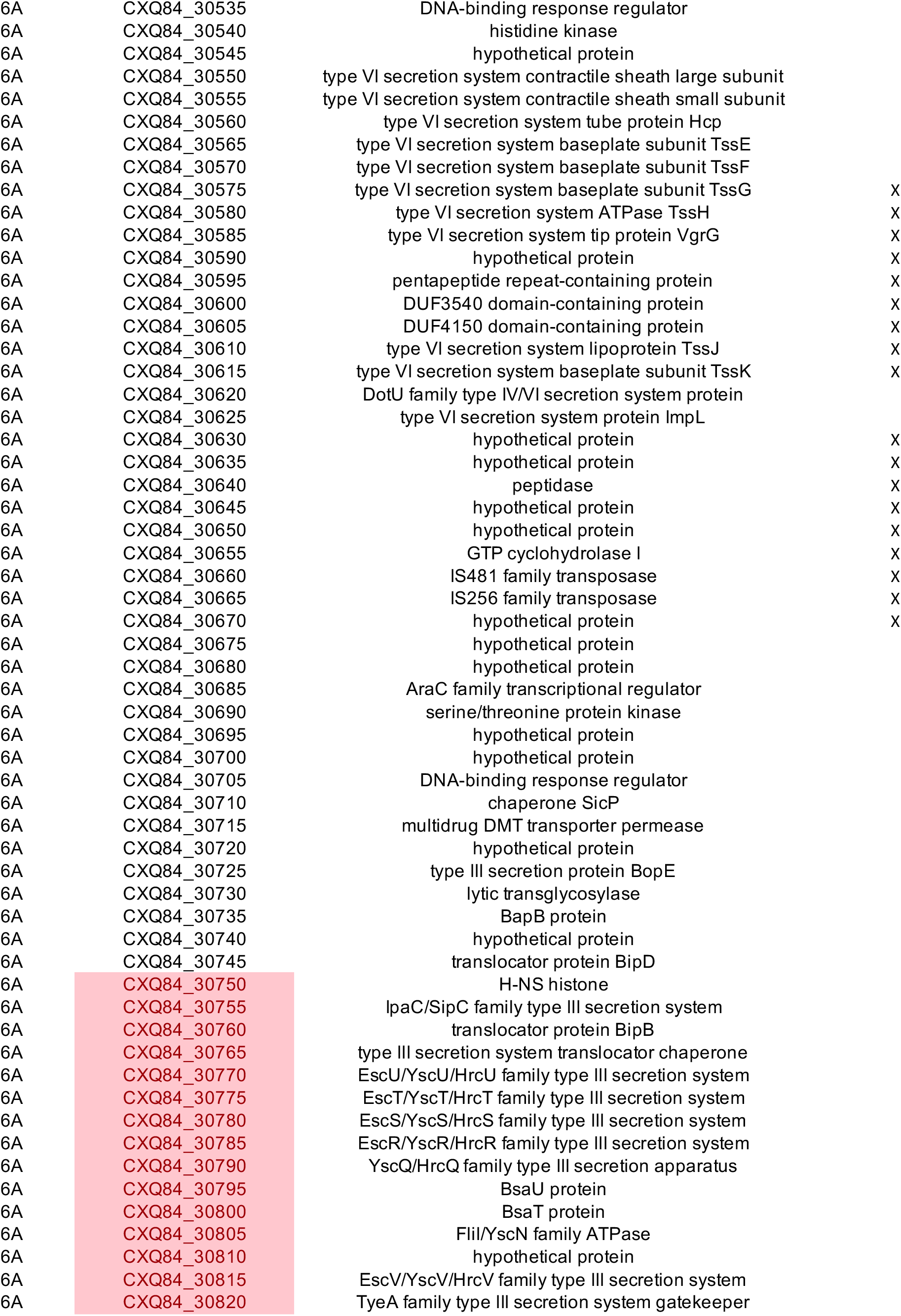
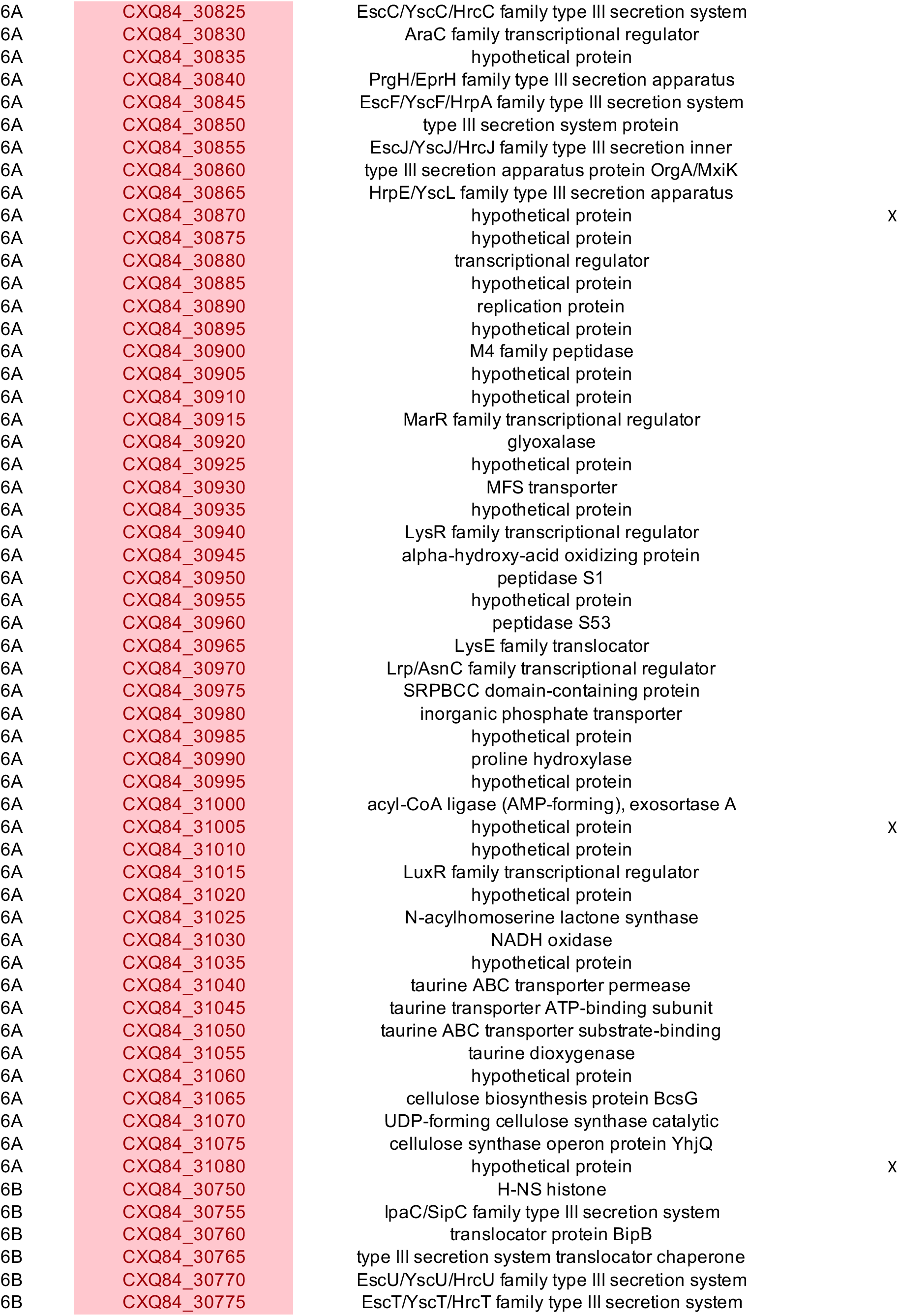
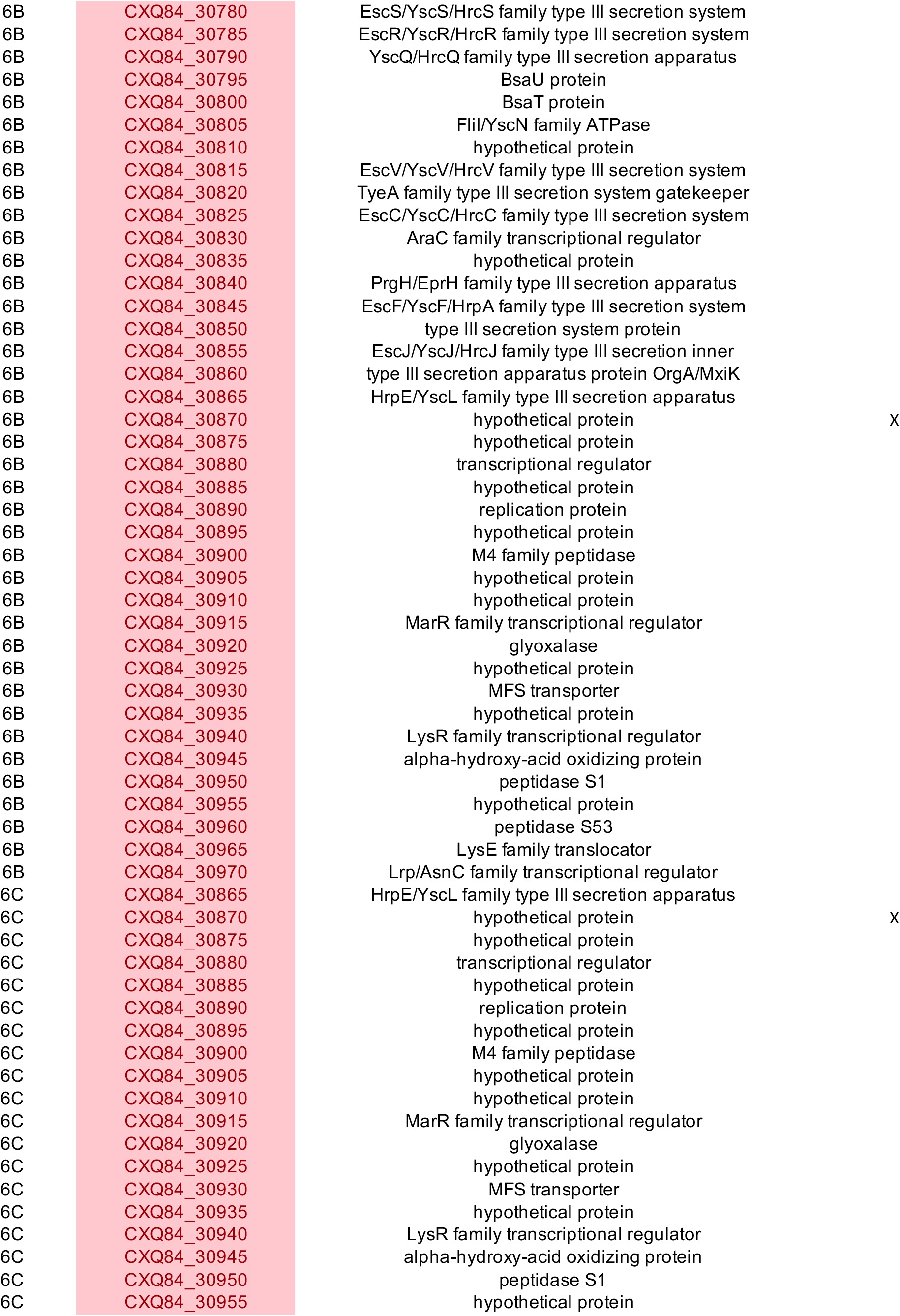
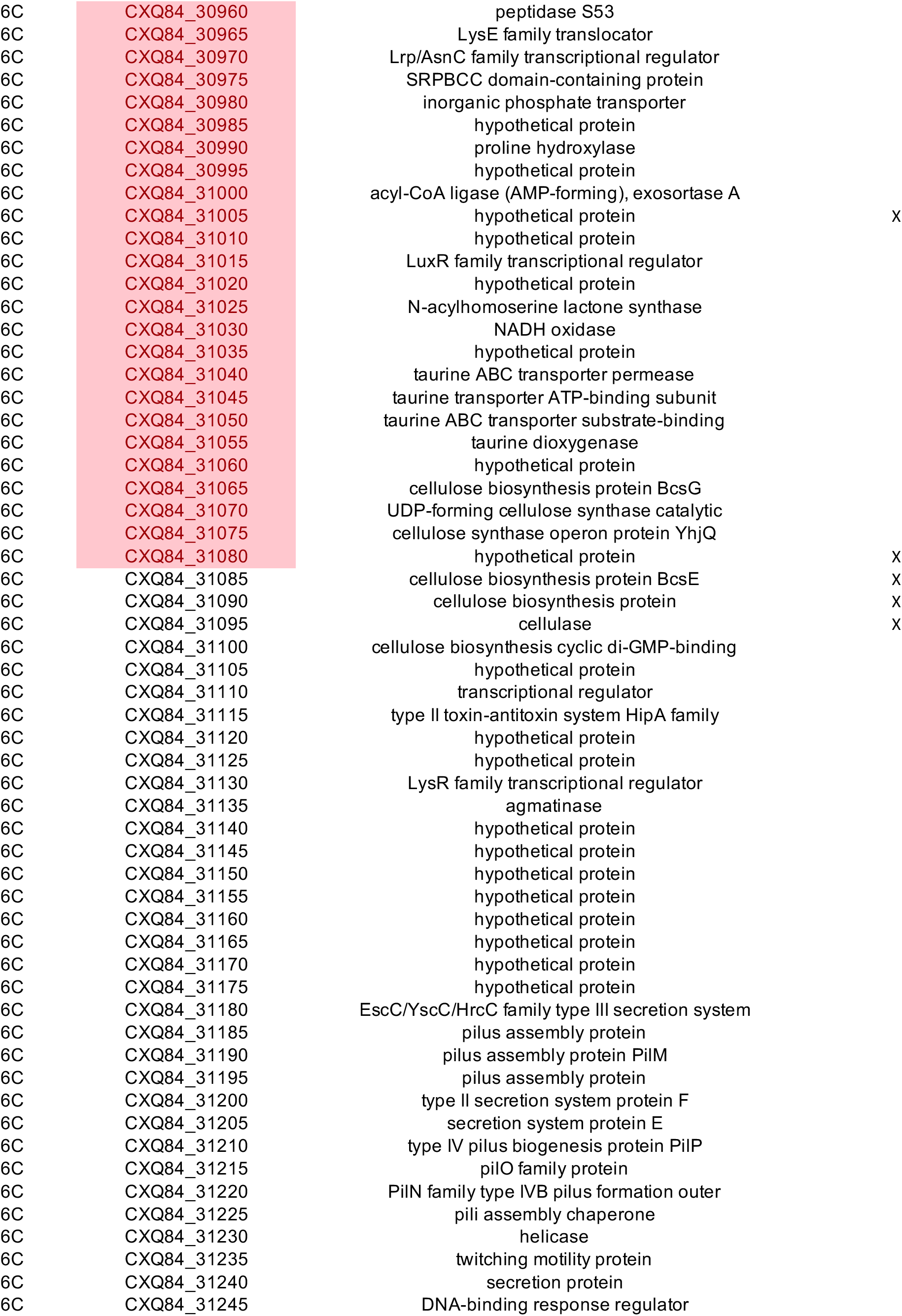
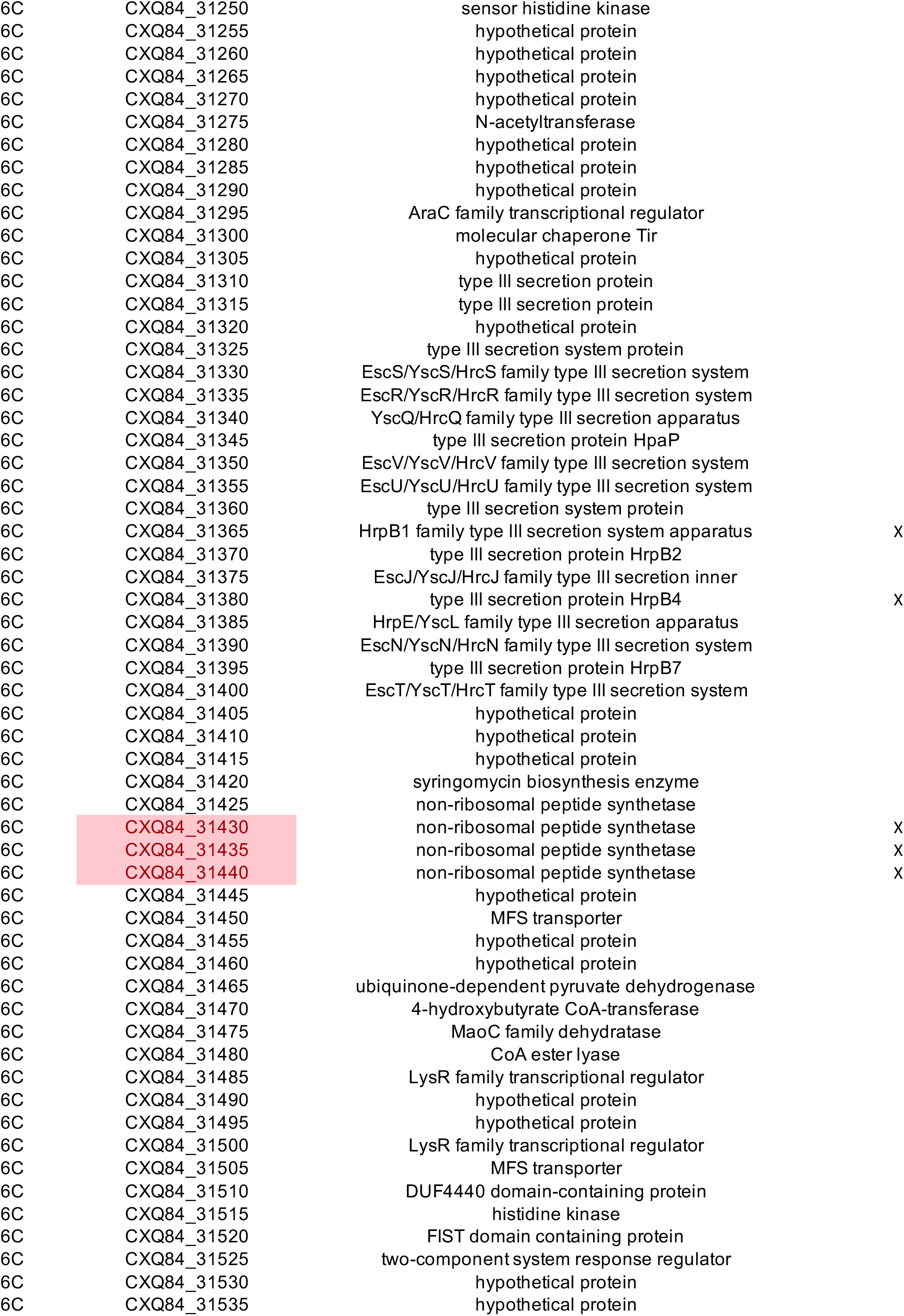
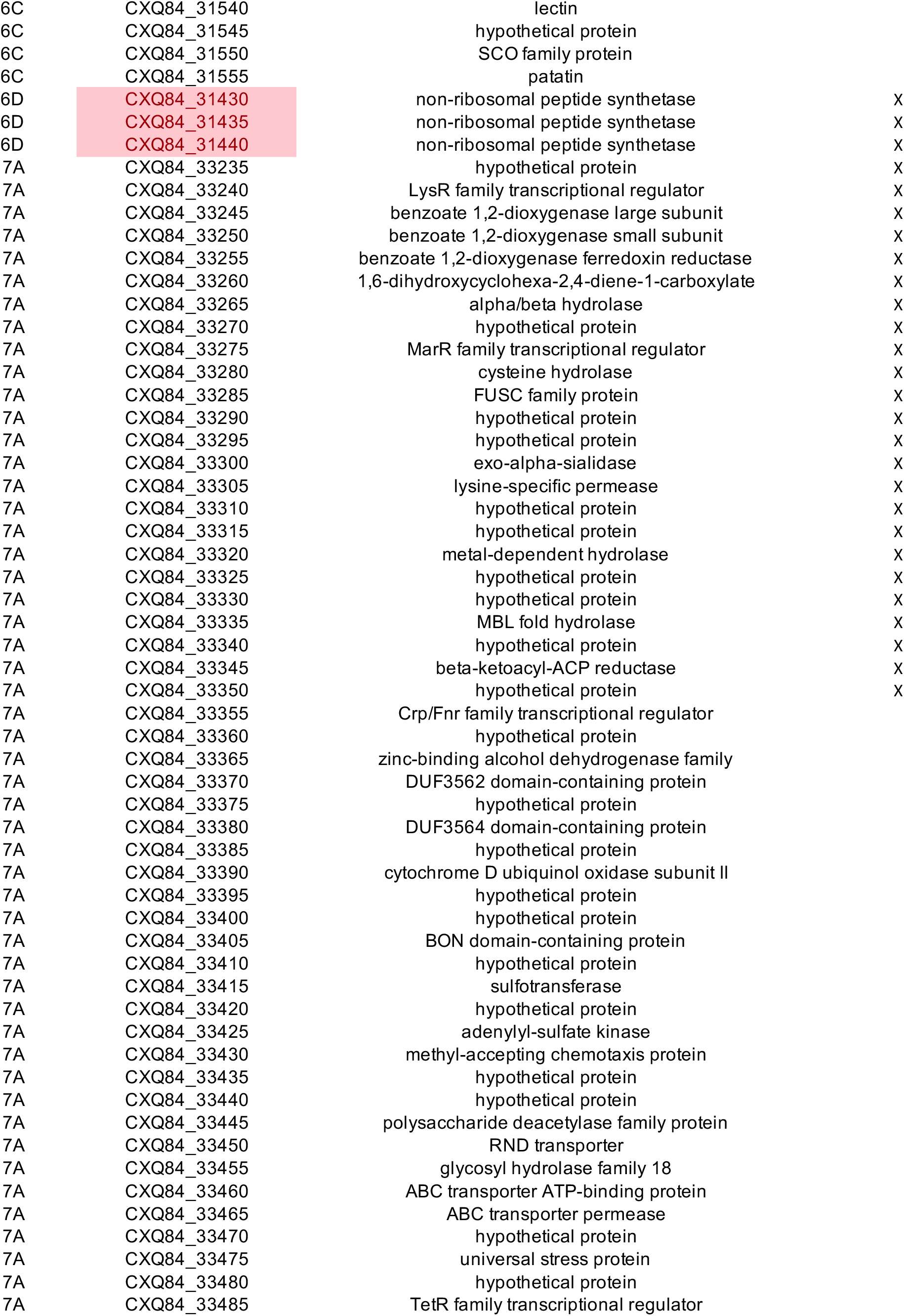
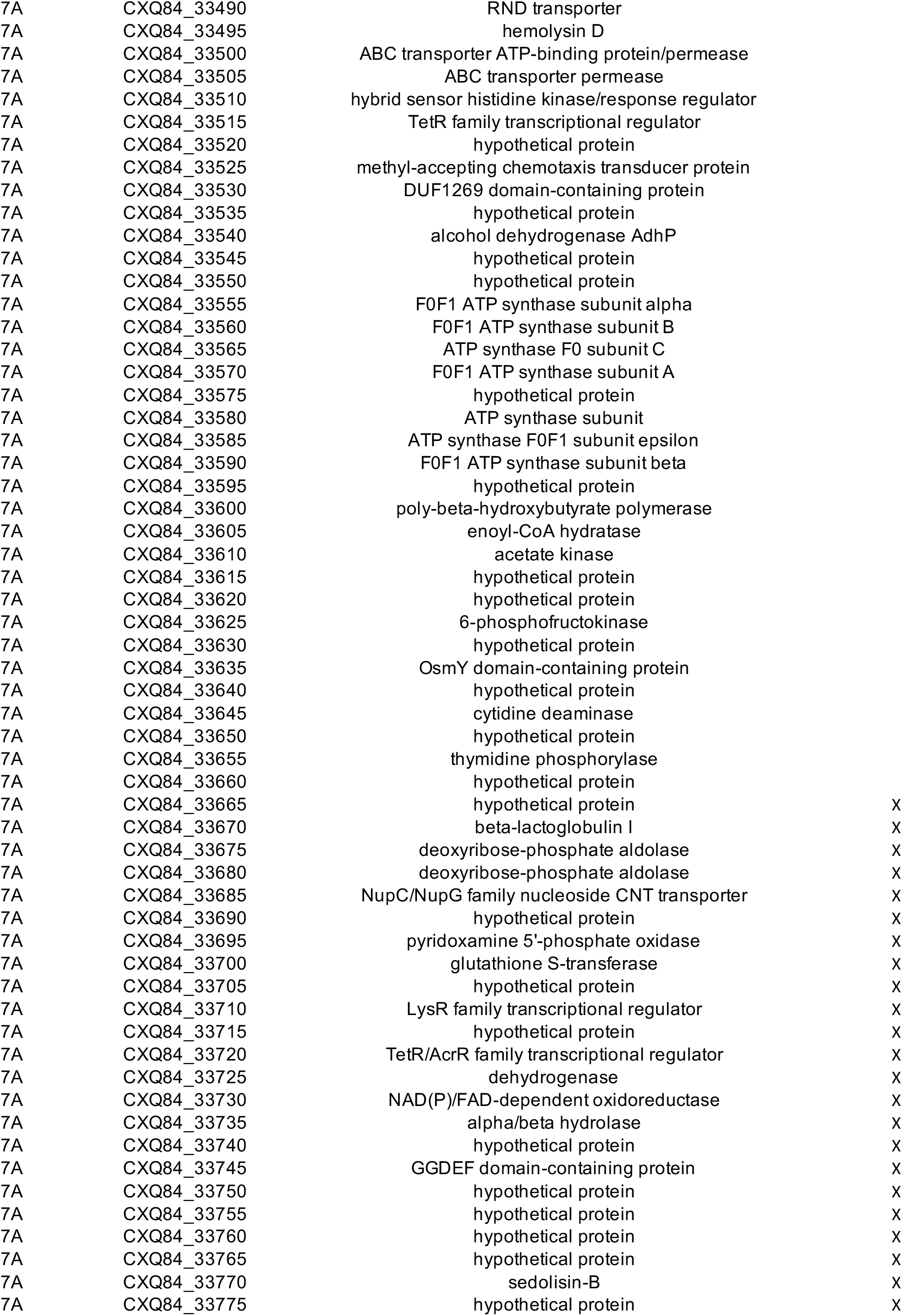
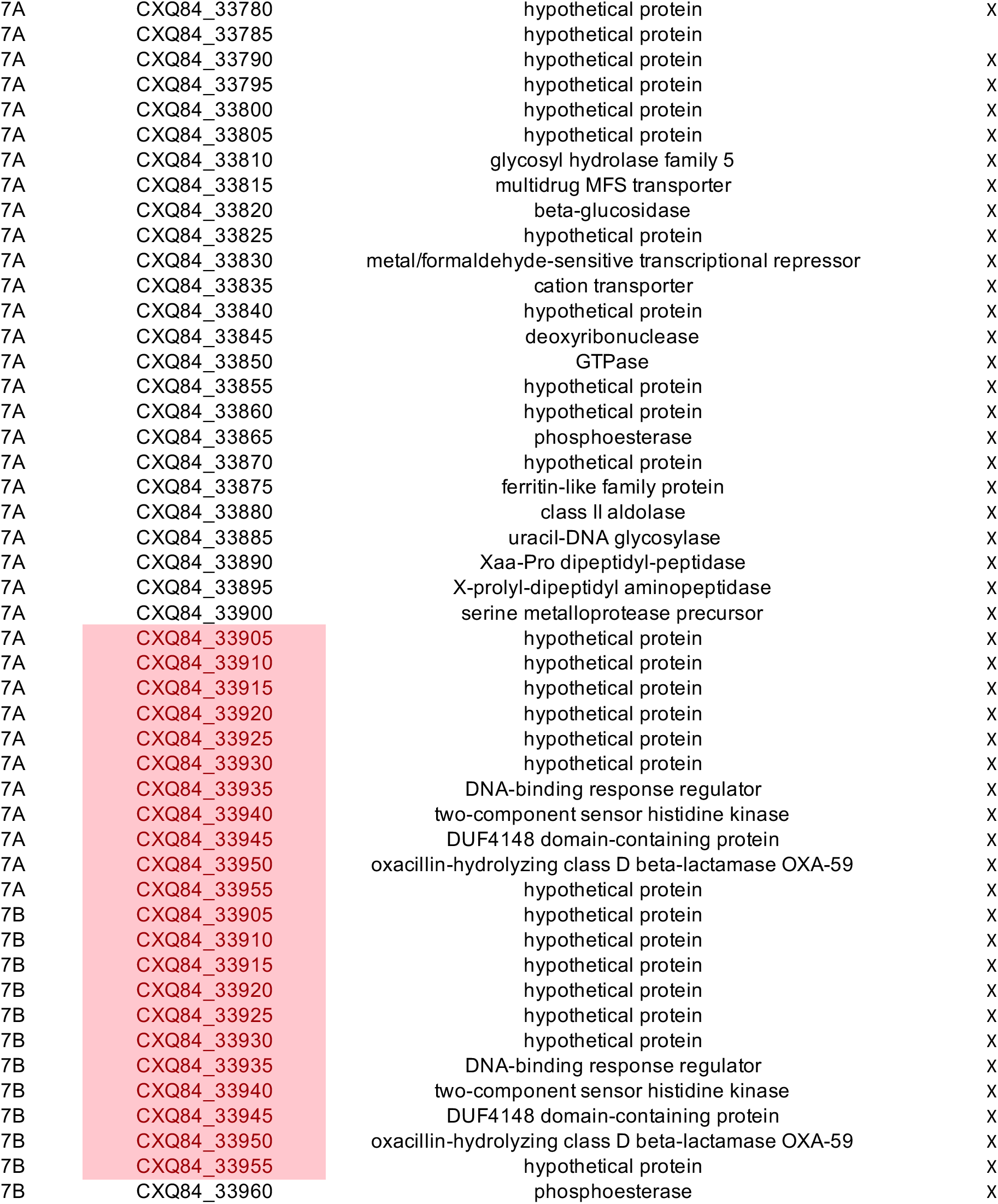
Details of large deletions.

**S4:** Generalized stepping stone analysis.

|  | Constant | Exponential | Skygrid | Skyride |
| --- | --- | --- | --- | --- |
| Strict | -1132.98 | -1139.11 | -1125.51 | -1453.11 |
| Lognormal | -1132.11 | -1139.56 | -1127.32 | -1463.60 |

**S5 Table:**
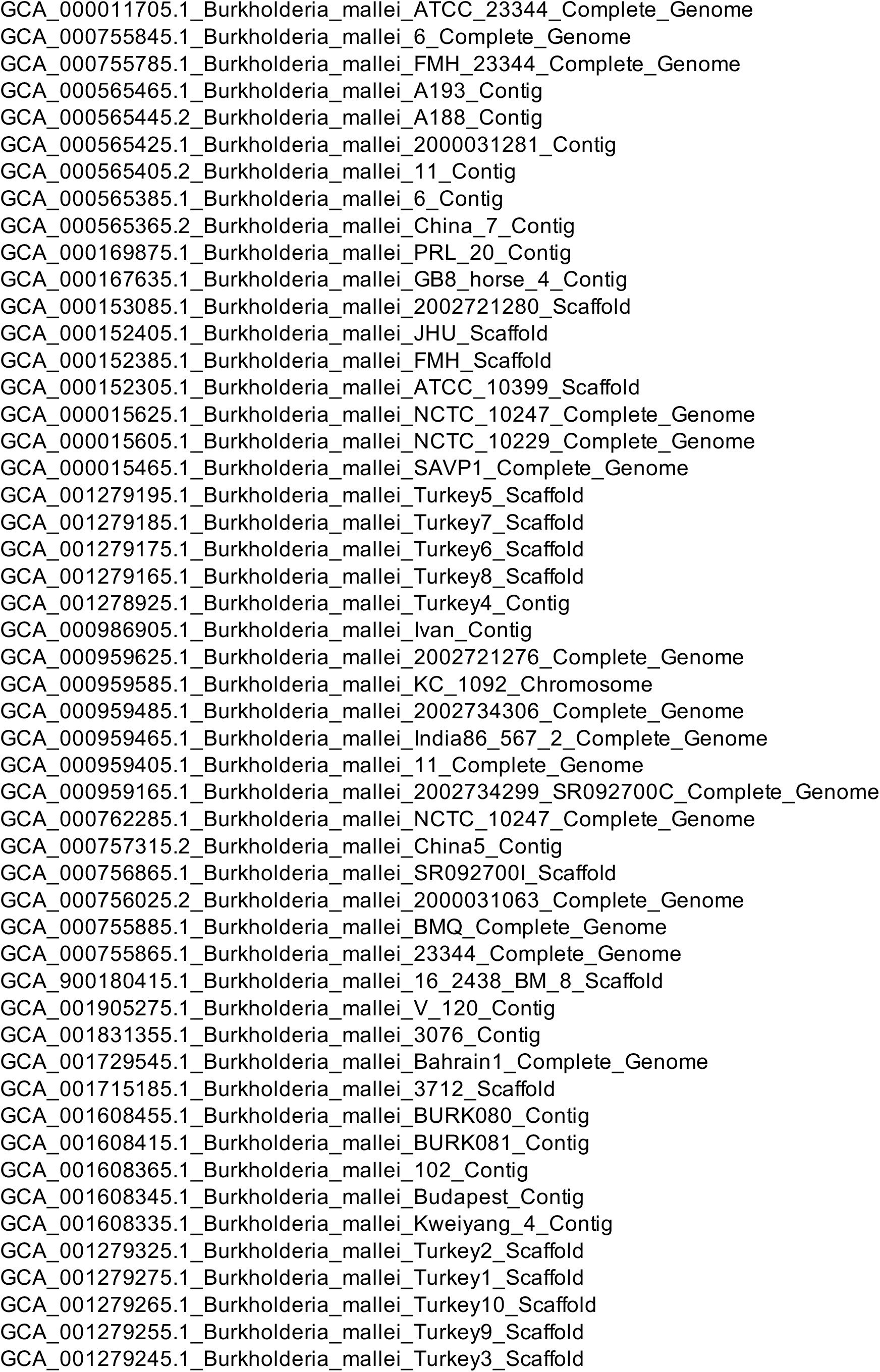
*B. mallei* accession numbers.

**S6 Table:**
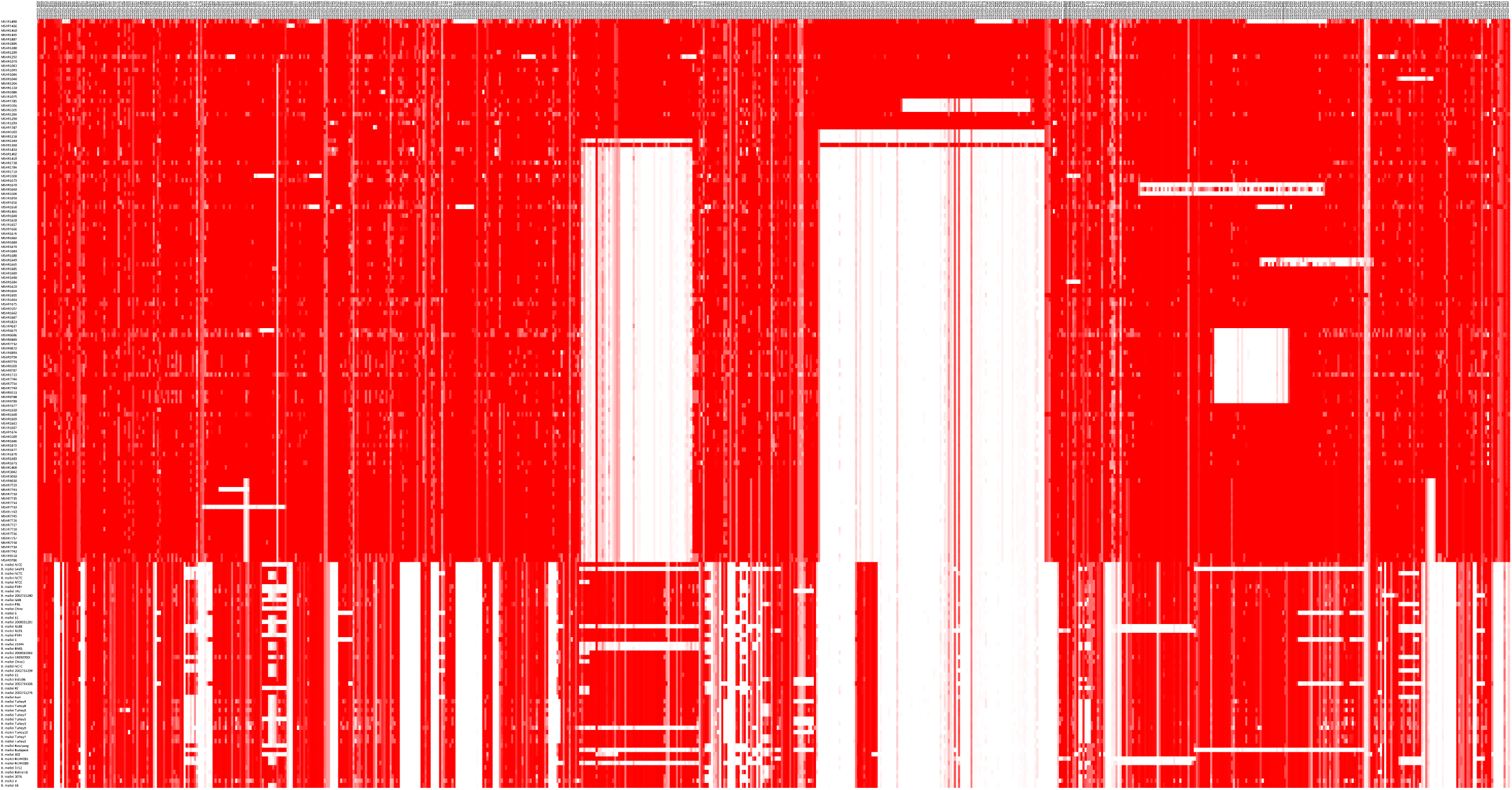
BSR.

**S7 Table:**
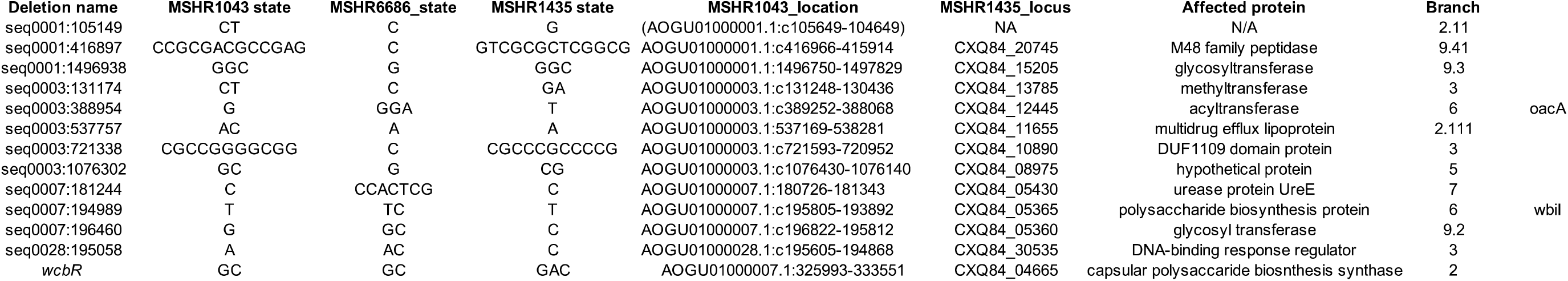
Small deletions.

**S8 Table:**
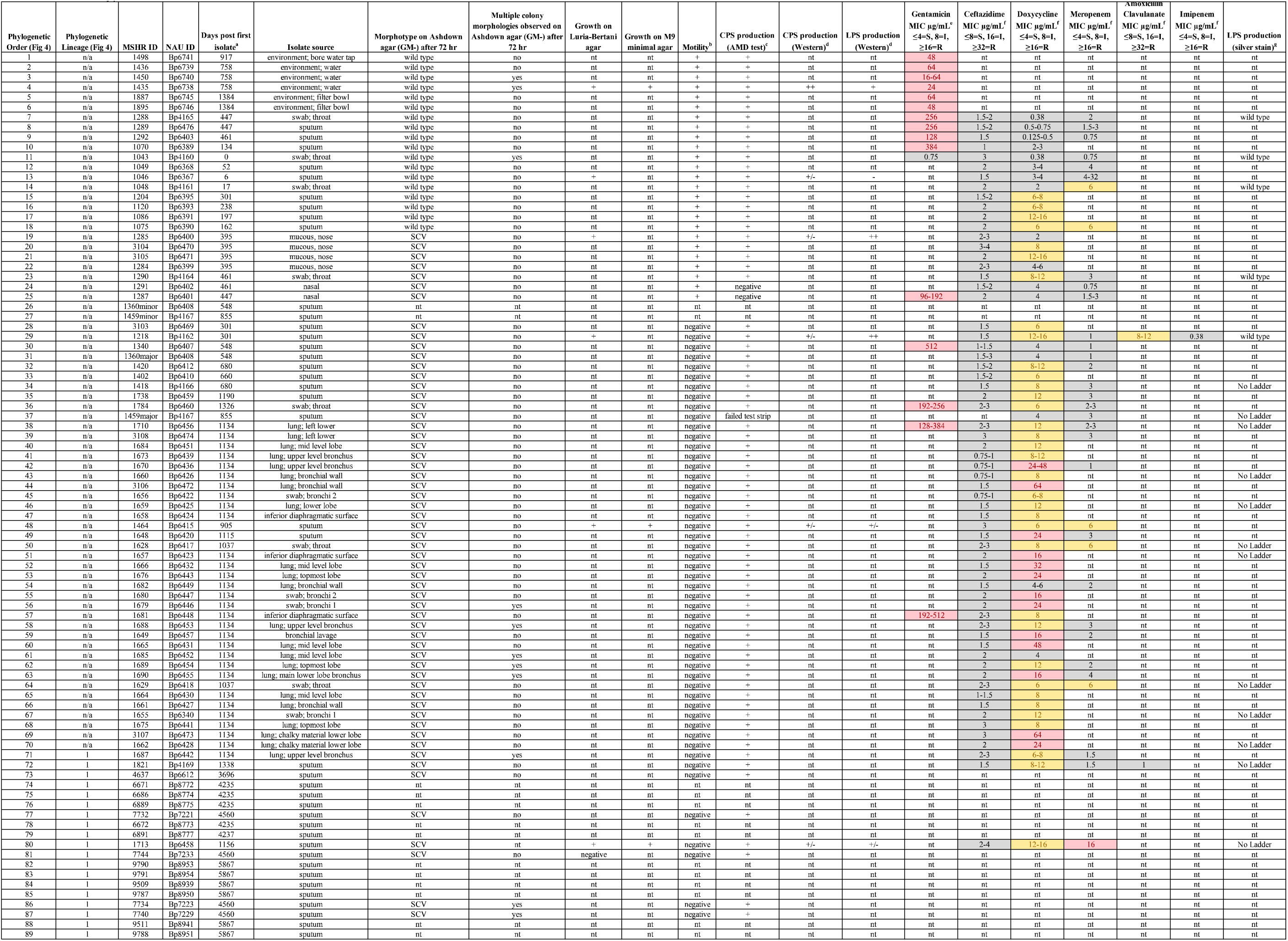
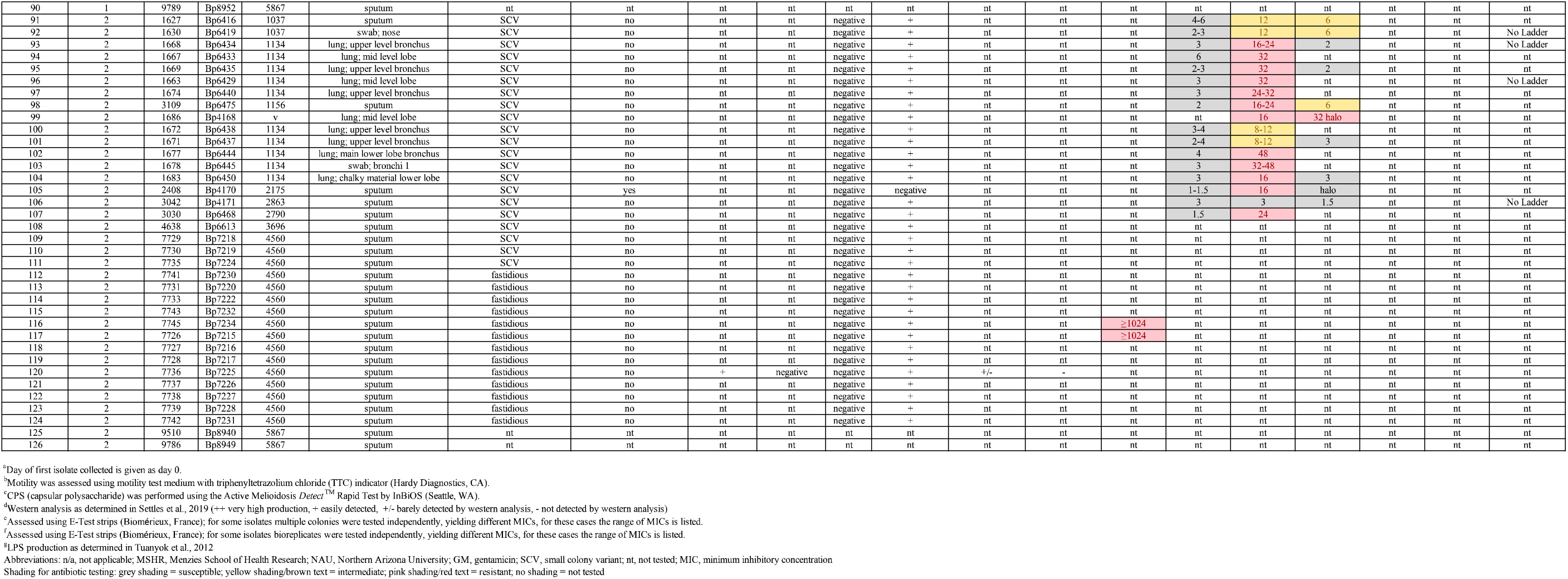
Phenotype data.

**S9 Table:**
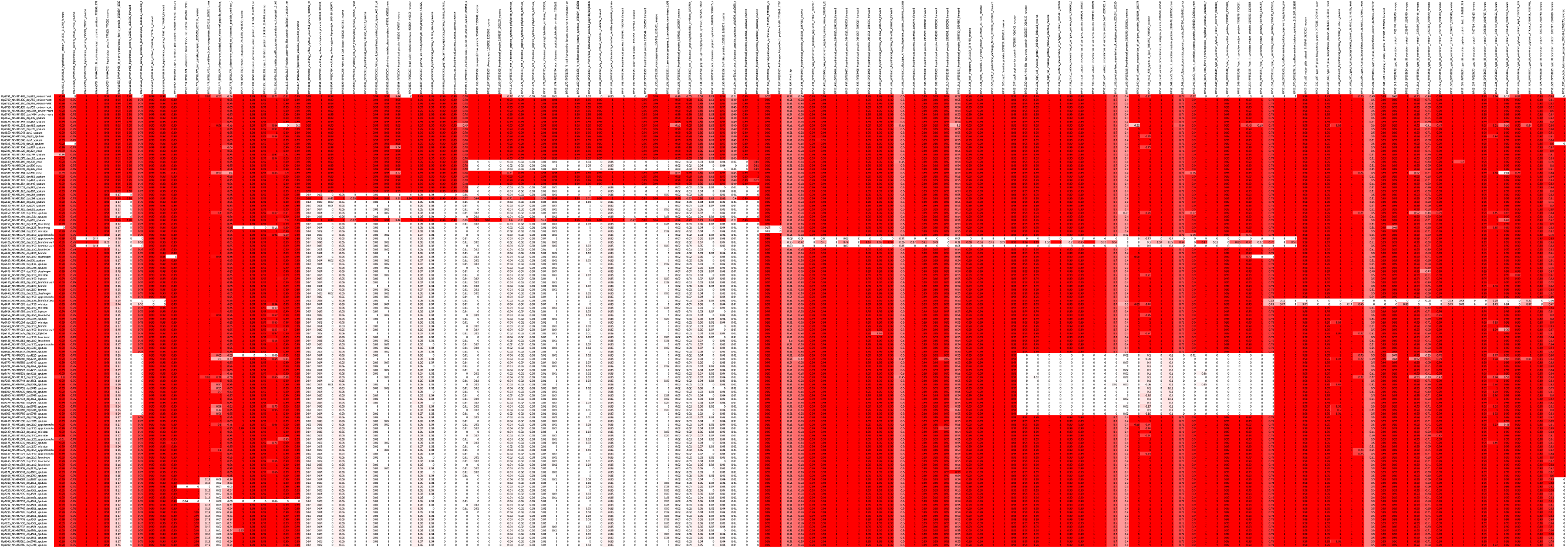
Virulence gene screen.

## Acknowledgments

We would like to thank Vanessa Rigas for assistance with culturing and Patient 314 for supporting this study over the past 18 years by being involved in our research. This work was funded under the State of Arizona Technology and Research Initiative Fund (TRIF), administered by the Arizona Board of Regents, through Northern Arizona University. This study was supported by grants from the Australian National Health and Medical Research Council: grant numbers 1046812, 1098337 and 1131932 (The HOT NORTH initiative).

